# Intricate Dynamical Cross-Talk Between p53 Protein and Cell Cycle Regulators Governs Mammalian Cell Fate

**DOI:** 10.64898/2026.06.07.730771

**Authors:** Kajal Charan, Sandip Kar

## Abstract

In mammalian cells, under normal circumstances, the p53 protein exhibits oscillatory dynamics in response to DNA damage and maintains the cells in a cell-cycle-arrested state. Intriguingly, some cells can escape this cell-cycle-arrested state even after prolonged DNA damage, and often undergo mitotic catastrophe. In this context, the precise role of p53 dynamics and its complex interplay with cell-cycle regulation remain poorly understood. Herein, by constructing a comprehensive network model, we have identified crucial crosstalk regulations between the p53 protein and key cell-cycle regulators that enable some cells to escape cell-cycle arrest during prolonged DNA damage. The model further illustrates a probable cellular mechanism underlying mitotic catastrophe and predicts ways to induce it in a therapeutically relevant manner.

## Introduction

Mammalian cells respond to DNA damage by halting the cell cycle and allowing DNA repair to maintain genomic integrity (Fulda *et al*, 2010; Abuetabh *et al*, 2022). One such important regulator of this process is p53, which activates and transmits the DNA damage signal to the cell cycle machinery (Levine, 1997). It is believed that the DNA damage-mediated dynamic regulation of p53 determines cell fate decisions (Purvis *et al*, 2012). Specifically, oscillations in p53 expression are associated with cell-cycle arrest, while sustained high levels lead to apoptosis (Purvis *et al*, 2012; Yang *et al*, 2018; Stewart-Ornstein & Lahav, 2017). However, recent findings suggest that not only p53 dynamics but also its interactions with cell cycle regulators determine cell fate, especially when cells are observed over longer periods of DNA damage (Reyes *et al*, 2018; Tsabar *et al*, 2020). For example, Reyes et al. showed, through long-term live-cell imaging, that some cells can escape cell cycle arrest and continue dividing even in the presence of DNA damage(Tran *et al*, 2023; Reyes *et al*, 2018). This escape may lead to genomic instability, which often manifests as micronuclei and is frequently associated with cancer (Fujimaki *et al*, 2025). This raises an important question: Is this escape solely driven by p53 dynamics, or does it result from a more complex set of interactions between key cell-cycle regulators and p53 signalling? If so, identifying these interactions and key cell cycle regulators can be extremely important from a therapeutic perspective, as it can help eliminate aberrant cells that escape cell cycle arrest during DNA damage.

Moreover, these escaping cells can even undergo mitotic catastrophe (Tsabar *et al*, 2020). Tsabar et al. have observed that cells that escape cell-cycle arrest exhibit a post-mitotic switch in p53 dynamics (Tsabar *et al*, 2020), which was associated with mitotic catastrophe. Normally, the mitotic catastrophe is considered a prelude to apoptosis or necrosis and is often associated with aneuploidy(Baptiste-Okoh *et al*, 2008; Sazonova *et al*, 2021; Mc Gee, 2015; Vakifahmetoglu *et al*, 2008). This might serve as a general mechanism for preventing cells with faulty genetic status from forming. However, how this process relates to p53 dynamics induced by DNA damage remains unclear. Interestingly, the mitotic catastrophe is observed during many cancer treatments, yet the underlying mechanisms remain poorly defined (Haschka *et al*, 2018; Bai *et al*, 2023; da Costa *et al*, 2023). Collectively, these findings suggest that cross-talk regulation between cell cycle regulators and p53 is extremely crucial for understanding p53-mediated cell fate decision-making. However, the mechanistic understanding of this cross-talk regulation is limited and requires further investigation.

Mathematical modelling is a powerful tool for understanding the mechanisms underlying such complex phenomena. Earlier modelling attempts that combined p53 and cell cycle regulation were mostly deterministic models or focused primarily on capturing cell cycle arrest (Jonak *et al*, 2016; Jung *et al*, 2021; Zhang *et al*, 2010; Heldt *et al*, 2018; Chong *et al*, 2015; Tian *et al*, 2017). The role of stochastic fluctuations (intrinsic fluctuations due to low copy numbers of mRNA’s or proteins in the concerned network) in modulating cell cycle regulation and facilitating the escape has not been explored. We believe that this can be of great importance, as even under DNA damage conditions, only a fraction of cells escape the therapy, others arrest the cell division cycle. Thus, in this article, we coupled our previously proposed p53 regulatory network model with an established cell-cycle regulatory network (Charan *et al*, 2022; Govindaraj *et al*, 2022) and introduced stochastic fluctuations into the coupled model. Our model simulations demonstrate that some fraction of cells escape cell-cycle arrest during prolonged DNA damage. Importantly, the model allowed us to identify which cell-cycle regulators are preferentially modulated to facilitate this escape from the cell-cycle-arrested state. The mechanistic insight from our model also aligns with relevant experimental studies (Yang *et al*, 2017,Segeren *et al*, 2020)). Further, we propose a mechanism underlying mitotic catastrophe and reconcile the corresponding dynamical features of various proteins, including p53 (Tsabar *et al*, 2020). Our model also predicts the probable ways in which mitotic catastrophe can be induced and modulated. In summary, this article illustrates how the dynamic crosstalk between p53 and cell-cycle regulators influences cell fate under DNA-damage conditions.

### Model

We built a coupled model of p53 and cell cycle regulation by adapting previously established models of p53 and cell cycle regulatory networks (Charan *et al*, 2022; Govindaraj *et al*, 2022). The minimal p53 signaling network (**Module 2**) incorporates the delayed p53–Mdm2 negative feedback loop, with the delay introduced by different p53 oligomers (Gaglia *et al*, 2013; Gaglia & Lahav, 2014). Here, the activation of Mdm2 is facilitated by distinct p53 oligomers (Barak *et al*, 1993; Fischer *et al*, 2016), and once Mdm2 is phosphorylated to its active form, it can degrade p53 (Haupt *et al*, 1997). The extent of Mdm2-mediated degradation varies across oligomers, with the highest for the tetramer and the lowest for the monomer (Gaglia *et al*, 2013; Gaglia & Lahav, 2014). The model further incorporates the dynamics of DNA damage generation and repair. DNA damage facilitates the degradation of Mdm2, thereby mimicking ATM-mediated phosphorylation and degradation of Mdm2, which later stabilises p53 (Charan *et al*, 2022; Khosravi *et al*, 1999) (**Fig. 1)**.

**Fig. 1.**
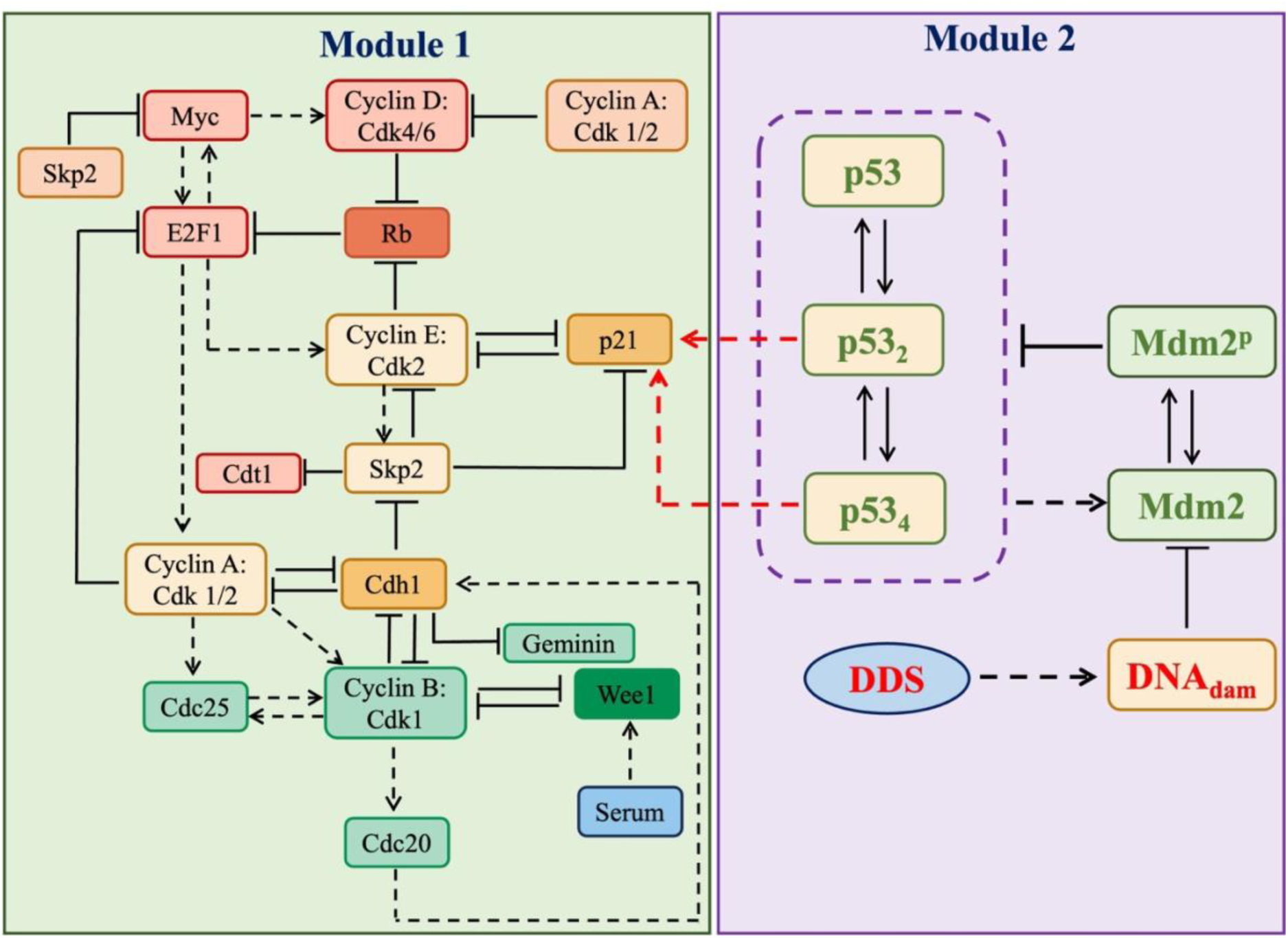
The proposed coupled cell cycle-p53 regulatory network. **Module 1** and **Module 2** represent the cell cycle and p53 regulatory networks, respectively. (Dashed and solid arrows denote activation and biochemical reactions, respectively. The solid hammer-headed lines depict inhibition.)

The cell cycle regulatory network (**Module 1**) used in this study is adapted from Govindaraj et al. (Govindaraj *et al*, 2022). This model is a serum-dependent cell cycle network that captures the dynamics of different cyclins and other cell cycle regulators during cell cycle progression (Govindaraj *et al*, 2022) (**Fig. 1)**. In this model, serum activates Cyclin D (CycD) and Myc. CycD initiates partial phosphorylation of Rb, which begins to disrupt the Rb-E2F1 complex. However, the complete phosphorylation of Rb requires Cyclin E (CycE) activation. During the G_1_ phase, CycE activity is suppressed by p21, which forms a CycE-p21 complex and prevents CycE activation (Govindaraj *et al*, 2022). The partial E2F1 activity promotes CycE production. Once some free CycE is produced, it phosphorylates Skp2, protecting it from degradation by Cdh1. This creates positive feedback because Skp2 degrades p21, thereby making CycE fully active (Govindaraj *et al*, 2022) (**Fig. 1)**. At the same time, active E2F promotes transcription of Cyclin A (CycA). CycA phosphorylates and inactivates Cdh1, which further stabilises CycE and Skp2. Increased Skp2 promotes S-phase entry by inactivating Cdt1 (Govindaraj *et al*, 2022) (**Fig. 1)**.

After S-phase entry, CycA activates CycB to facilitate mitosis and exit from the cell cycle. CycA also activates Cdc25 transcription, although it must be phosphorylated by CycB to become active. CycB begins to accumulate during the G_2_ phase and forms positive feedback loops with Cdc25 and Wee1. Once CycB becomes fully active, mitosis begins (Govindaraj *et al*, 2022) (**Fig. 1)**. During mitosis, CycB activates Cdc20, which, in turn, activates Cdh1. At the end of mitosis, Cdh1 is activated and reduces active CycB levels by phosphorylation. This sudden drop in CycB levels leads to cell division (Govindaraj *et al*, 2022; Tyson & Novak, 2001). Experimentally, cell cycle progression can be monitored following geminin expression using single-cell reporter constructs (Reyes *et al*, 2018). The inactivation of geminin by Cdh1 causes geminin expression to rise during entry into the S phase and drop during cell division (Govindaraj *et al*, 2022; Sakaue-Sawano *et al*, 2008) (**Fig. 1)**.

During the normal cell cycle, cells pass through several checkpoints that monitor internal and external conditions. Importantly, the G_1_ checkpoint inspects for DNA damage (Hyun & Jang, 2015). If damage is detected, the cell activates the cell cycle inhibitor p21 and it halts the cell cycle in the G_1_ phase (Fischer *et al*, 2016) (**Fig. 1)**. Experimental evidence shows that p21 is activated by p53 (Reyes *et al*, 2018; Fischer *et al*, 2016). In particular, p53, in its dimeric and tetrameric forms, promotes and activates p21 transcription (Gaglia *et al*, 2013). To link DNA damage effects to cell cycle progression, we integrated these additional interactions to establish a connection between the p53 signalling and cell cycle regulatory networks (Gutu *et al*, 2023; Phillips *et al*, 2019). We converted these network interactions into ordinary differential equations and simulated them using XPPAUT (https://sites.pitt.edu/phase/bard/bardware/xpp/xpp.html). To capture the inherent stochasticity of biological systems, we treated all interactions as independent reactions and simulated them using Gillespie’s stochastic simulation algorithm (Gillespie, 1976). The related parameter values and their descriptions are provided in **Tables S1-S8**.

## Results and Discussion

### Deterministic simulation demonstrates cell cycle arrest in the presence of DNA damage

First, we employed the coupled deterministic p53-cell cycle model and examined its response in the presence and absence of DNA damage. Under normal conditions (without DNA damage), p53 maintains a low basal level (Loewer *et al*, 2013). As a result, p21 exhibits typical dynamics: it rises during the G_1_ phase and drops during the G_1_-S transition due to CycE activation. This allows normal cell cycle progression (Reyes *et al*, 2018; Govindaraj *et al*, 2022). In contrast, p53 is activated and oscillates under DNA-damage conditions. This activation leads to sustained p21 expression (Reyes *et al*, 2018). Elevated p21 suppresses CycE activity and prevents the G_1_-S transition by halting the cell cycle at the G_1_ phase (Karimian *et al*, 2016).

Our deterministic simulations (**Fig. 2A-C)** demonstrate that, upon DNA damage, oscillatory p53 activates p21 (Fischer *et al*, 2016). Consequently, p21 inhibits CycE expression, effectively arresting the cell cycle. As DNA damage is gradually repaired (∼80 hr in **Fig. 2A**), the relative expression of p21 and p53 declines, allowing the normal cell cycle to resume (Barr *et al*, 2016), as is evident from the CycB oscillatory expression (**Fig. 2D)**. Through this model, the DNA damage-induced cell cycle arrest was successfully captured (**Fig. 2A-D).** However, this model is completely deterministic and cannot explain the intermittent cell-cycle escape phenomenon. Thus, we next investigated whether fluctuations in the coupled p53-cell-cycle network can modulate cell-cycle arrest during prolonged DNA damage.

**Fig. 2.**
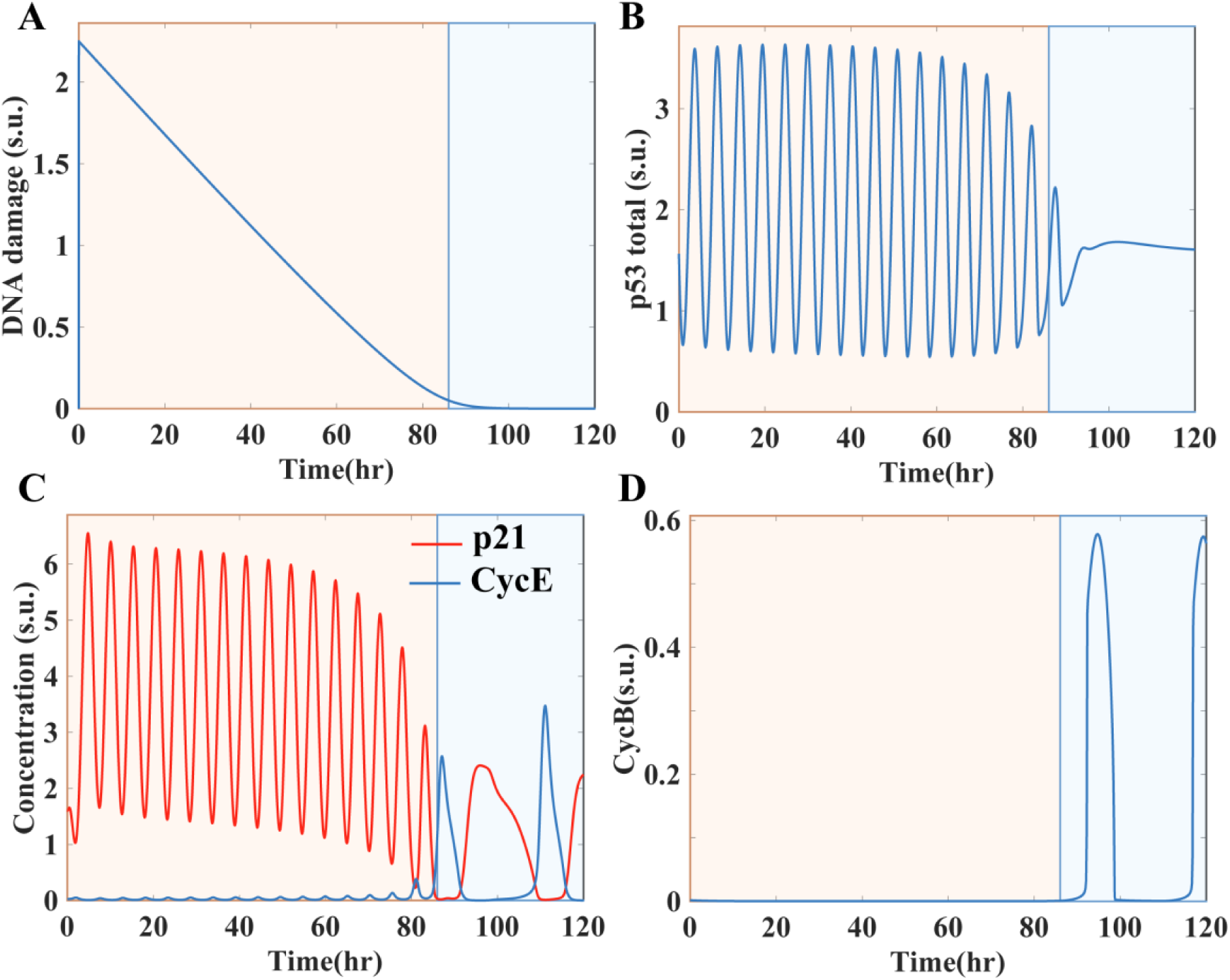
Deterministic simulation of coupled cell cycle p53 network. Time profiles of **(A)** DNA damage, **(B)** p53 total, **(C)** CycE and p21, **(D)** CycB, showing the DNA damage-induced cell cycle arrest.

### Stochastic fluctuations drive cell cycle arrest escape in a subset of cells

We incorporated stochastic fluctuations into the coupled network using Gillespie’s stochastic simulation algorithm (Gillespie, 1977). We first simulated cells under normal conditions without external DNA damage (DDS = 0). In our model, intrinsic DNA damage is randomly generated, consistent with experimental findings that low levels of endogenous DNA damage occur even during normal cell cycle progression (Loewer *et al*, 2013). This intrinsic DNA damage induces random p53 pulses **(Fig. 3A (i))**, but these are insufficient to arrest the cell cycle **(Fig. 3A (ii))**. As a result, cells continue to divide normally, as indicated by Geminin expression **(Fig. 3A(iii))** before mitosis.

**Fig. 3.**
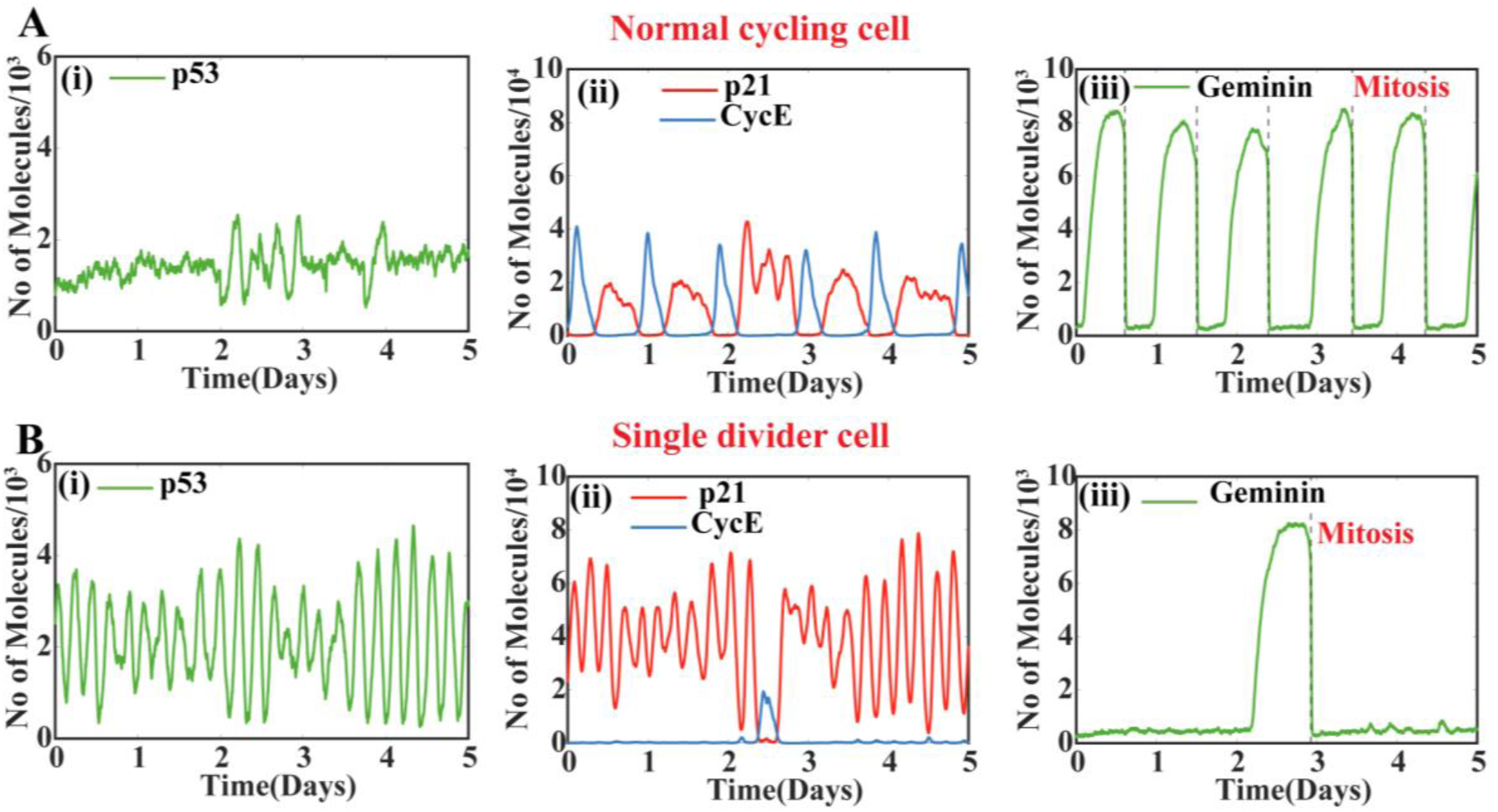
Stochastic time profiles for p53 and cell cycle regulators. Representative Time profile of **(i)** p53, **(ii)** p21 and CycE, and **(iii)** Geminin for **(A)** the normal cycling cell and **(B)** the single divider cell during DNA damage.

To assess the effect of external DNA damage, we applied a transient DNA damage signal (DDS = 30 a. u.) for 2 minutes, then monitored the system for 5 days with DDS = 0. Under these conditions, most cells remain in a cell-cycle arrest and do not divide, as confirmed by CycB and Geminin levels **(Fig. S1B)**. However, a few cells escape cell-cycle arrest and divide. Cell division is marked by a peak in CycB, which signals the exit from mitosis (**Fig. 3B)**. In the time profile of these escaping cells, we observed a sharp transition between CycE and p21 levels (**Fig. 3B(ii))**, suggesting a switch-like activation of CycE that enables escape from arrest while p53 maintains the oscillatory dynamics (**Fig. 3B(i))**. These results are consistent with the experimental observations by Reyes et al., in which a subset of cells escapes from cell-cycle arrest despite DNA damage (Reyes *et al*, 2018).

### The escaping tendency of cells varies with DNA damage dosage

To explore how this escape depends on different DNA damage doses, we simulated varied extent of DNA damage by altering DDS values from 0 to 200 a.u.. Our simulation demonstrates that as the DNA damage dose increases, p53 expression increases, leading to higher p21 expression and a lower probability of cells escaping cell-cycle arrest **(Fig. 4A)**.

**Fig. 4.**
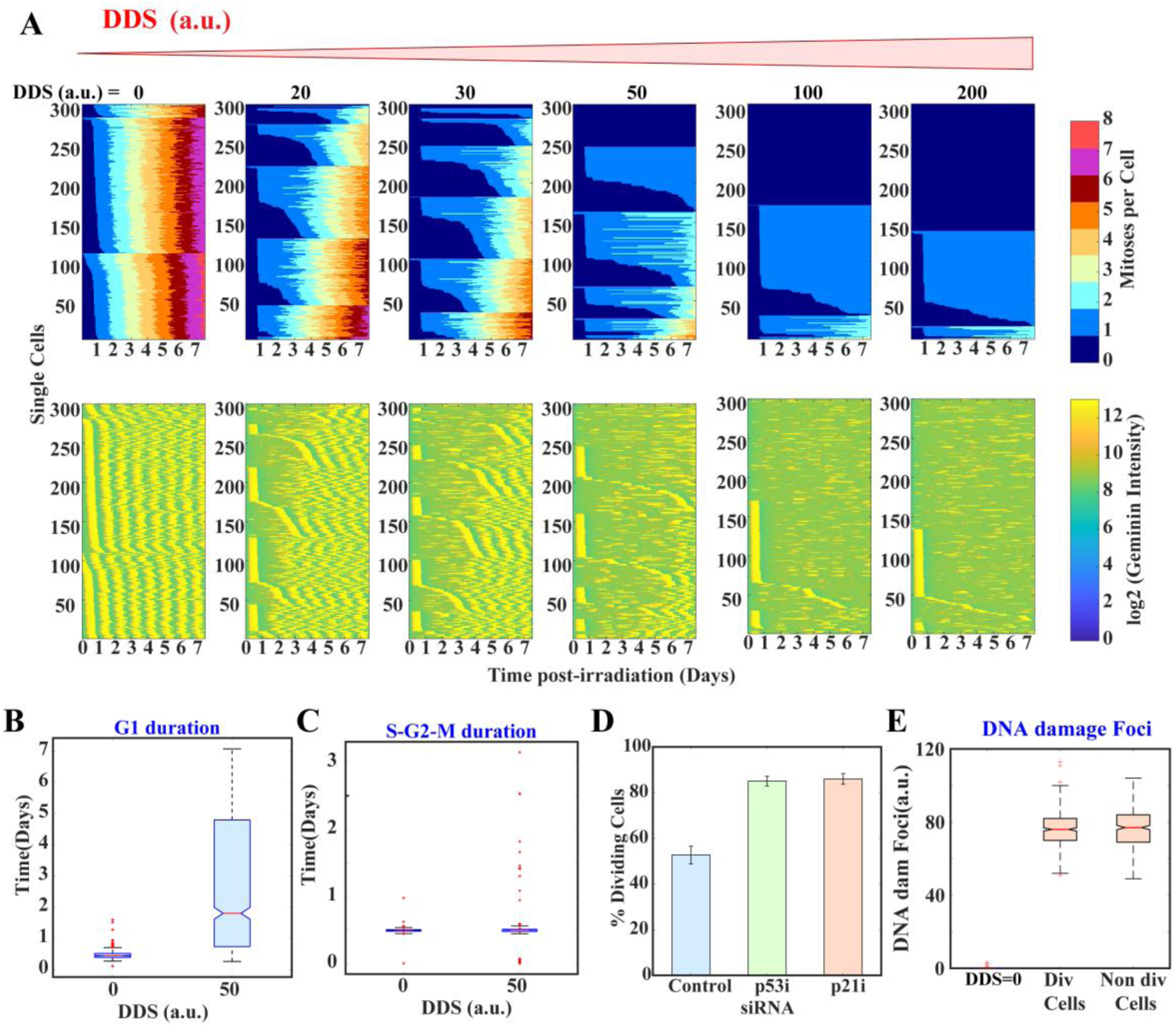
DNA damage-mediated cell cycle arrest during G1 phase. **(A)** Division dynamics in terms of mitoses per cell (top) and Geminin expression (bottom) profiles at different DNA damage signal doses during a 7-day simulation (n=300 for each condition). **(B)** G_1_ and **(C)** S-G_2_-M duration at *DDS* = 0 and *DDS* = 50 a.u. DNA damage conditions. **(D)** p53 and p21 siRNA simulations showing an increased fraction of dividing cells (at *DDS* = 50 a.u.). **(E)** DNA damage foci number for *DDS* = 0, dividing cells, and non-dividing cells for *DDS* = 50 a.u. after 48 hours.

At DDS = 0, all cells underwent multiple rounds of division, and as the damage dose increased, the number of dividing cells decreased. At high doses, such as DDS = 100 a. u. or 200 a.u., most cells remained arrested, though a small fraction still divided (**Fig. 4A)**. Interestingly, some cells divided only once. Following the terminology of Reyes et al., we refer to these as single dividers (Reyes *et al*, 2018). Among them, cells that divide only after 48 hours of DNA damage are termed *single late dividers,* or *“Escapers”*. In the subsequent analysis, we have focused on these escaper cells to exclude cells that are already committed and divide immediately after DNA damage, as these escaper cells are important in the therapeutic context. To validate that DNA damage-induced cell cycle arrest occurs in the G_1_ phase, we quantified the G_1_ and S-G_2_-M durations of normal cycling cells (DDS = 0) and cells at DDS = 50 a. u. Our simulation shows that the G_1_ duration is significantly prolonged in DNA damaged cells (**Fig. 4B)**, whereas the S-G_2_-M duration remains unchanged (**Fig. 4C)**. This finding corroborates with experimental observations and confirms that cell cycle arrest occurs primarily in the G_1_ phase (Reyes *et al*, 2018).

Furthermore, to investigate whether cell cycle arrest or escape from the arrested state is driven by p53 and its downstream effector, p21, Reyes et al. silenced the p53 and p21 genes and observed an increased fraction of cells undergoing cell division, even in the presence of DNA damage (Reyes *et al*, 2018). Similarly, in our model simulations, reducing the transcription rates of p53 and p21 mRNAs (simulating the influence of selective siRNA) increases the fraction of dividing cells **(Fig. 4D)**. This further underscores the roles of p53 and p21 in mediating the DNA damage response. Finally, we examined whether DNA damage foci differ between non-divider and escaper cells **(Fig. 4E)**. Our simulation shows that DNA damage foci occur less frequently in the no DDS condition, as expected. However, there is no difference in foci levels between non-divider and escaper cells **(Fig. 4E)**. This suggests that escape is not due to reduced DNA damage.

### The model identifies the key cell cycle regulators influencing cell fate decision-making during DNA damage

So far, our stochastic model captures escape from cell-cycle arrest despite active DNA damage. However, an important question remains: is this escape driven solely by stochastic fluctuations, or do the inherent expression of specific cell-cycle regulators and their crosstalk with p53 dynamics also contribute to this phenomenon? Reyes et al. proposed that this escape is purely due to fluctuations in p53 amplitude (Reyes *et al*, 2018). In contrast, other studies suggest that basal or accumulated levels of signalling proteins and cell cycle regulators, such as MAPK (Yang *et al*, 2021), Cyclin-D (Yang *et al*, 2017), and E2F1 (Segeren *et al*, 2020), can also influence cell fate under DNA-damage conditions. To further investigate this important issue, we compared the inherent levels of an important cell cycle regulator, Geminin, in non-divider and escaper cells via stochastic simulations (**Fig. 5A**). Here, non-divider cells will never show a high Geminin peak; however, in escaper cells, either mitosis occurs, followed by a high Geminin peak, or the Geminin peak appears without a mitosis event. Interestingly, p53 expression did not differ significantly between escapers and non-dividers **(Fig. 5B(i))**. However, p21 levels were lower in single late-dividing (escaper) cells compared to non-dividers **(Fig. 5B(ii))**.

**Fig. 5.**
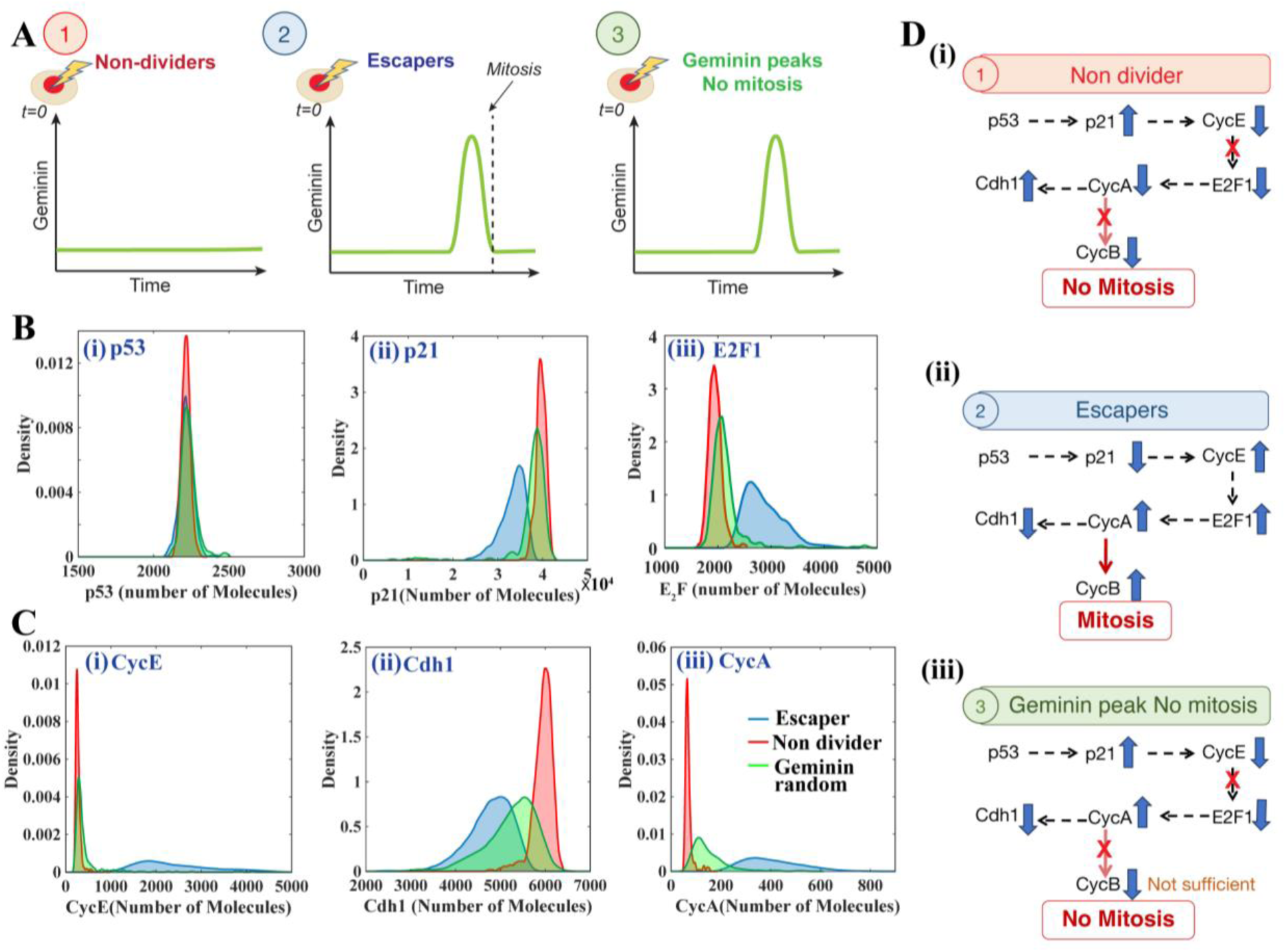
Predicting key cell cycle regulators during prolonged DNA damage condition. **(A)** Classification of cells into three categories under DNA damage condition: (1) Non-dividers without high Geminin, (2) Escaper cells undergoing mitosis after 48 hours of DNA damage, and (3) cells showing Geminin peaks without mitosis. **(B)** Expression profile of (i) p53, (ii) p21, and (iii) E2F1 in three cell categories. **(C)** Expression profile of (i) CycE, (ii) Cdh1, and (iii) CycA in three categories of cells. **(D)** Proposed mechanism underlying (i) non-dividers, (ii) escaper cells, and (iii) Geminin peaks without mitosis.

This shows that the p21 regulation by other cell cycle regulators also plays a major role in maintaining this DNA damage-mediated cell cycle arrest and can be responsible for escape. Thus, we analysed other upstream regulators of p21. We observed that the active form of E2F1 (i.e., DE in our model) accumulates in the escaper cells **(Fig. 5B(iii))**. It promotes CycE activation and, in turn, lowers p21 levels in escaper cells (**Fig. 5D(ii)**) in comparison to the non-divider cells (**Fig. 5D(i)**). These simulation results are consistent with existing experimental studies showing increased E2F1 activity in cells that progress through the cell cycle despite DNA damage (Segeren *et al*, 2020). Together, these results show that while p53 initiates cell cycle arrest through p21 activation, the maintenance of this arrested state depends on regulators of the broader cell cycle network (**Fig. 5D(i-ii)**).

### Random geminin peaks reveal failed attempts of division

Our simulations also capture occasional random peaks in Geminin in escaper cells without a mitosis event (as shown in **Fig. 5A**). Normally, Geminin levels rise during the S-G_2_-M phase and drop during mitosis. However, in some instances, we observed some random Geminin peaks that are not followed by mitosis (as indicated by CycB activation in our model). Experimental studies have supported the occurrence of similar events and described them as failed attempts at cell division (Tran *et al*, 2023; Zhang *et al*, 2023) and as resembling endocycle events (Fujimaki *et al*, 2025). To understand the mechanism behind these random Geminin peaks (**Fig. 5A**), we analysed the expression of other cell cycle regulators. We could not find any difference in expression of p53 and p21 in cells showing random Geminin peaks and non-divider cells (**Fig. 5B(i)-(ii)**). We found that active E2F1 levels in these cells are intermediate between those in non-dividers and those in escaper cells (**Fig. 5B(iii)**). This suggests partial E2F1 activation. As a result, CycA (**Fig. 5C(iii)**) become partially activated, but due to higher p21 levels, CycE cannot be released. However, elevated CycA levels can partially suppress Cdh1 (**Fig. 5C(ii)**), as evidenced by their expression levels. Although the cell attempts to progress through the cell cycle, the lack of CycE and CycA activation, and the intermediate levels of Cdh1, prevent CycB activation (**Fig.S1**). Because Cdh1 activity is not fully suppressed, the system resets back to G_1_ instead of cell cycle progression (**Fig. 5D(iii)**), making it look like an endocycle-like phenomenon (Fujimaki *et al*, 2025; Tran *et al*, 2023). The lack of mitosis module activation can also be confirmed by the expression levels of key mitotic regulators, such as Wee1 (**Fig. S2**), which show no significant differences between non-dividers and cells with random geminin peaks. Thus, our model analysis has provided a realistic, mechanistic interpretation of random Geminin peaks and dynamics in escaper cells under DNA damage conditions.

### Proposed mechanism of mitotic catastrophe in cells escaping cell cycle arrest

In the previous section, we examined how a subset of cells escapes DNA damage-induced G_1_ arrest. This raises a crucial question: what happens to these cells after they escape? Do they continue dividing normally, or is there a secondary mechanism that limits their proliferation? Tsabar et al. showed that p53 dynamics exhibit a sharp switch after mitosis in the presence of DNA damage, driven by mitotic catastrophe (Tsabar *et al*, 2020). They proposed that mitotic catastrophe is regulated by the PIDDosome complex, which is formed by PIDD and Caspase2 (Denisenko *et al*, 2016; Vakifahmetoglu *et al*, 2008; Tsabar *et al*, 2020). The DNA damage-induced activation of p53 upregulates the PIDD (p53-Induced Death Domain) protein, which binds to Caspase-2 (Casp2) to form the PIDDosome complex after defective mitosis. This complex inhibits Mdm2, thereby stabilising p53 and leading to a significant increase in p53 expression (Tsabar *et al*, 2020). However, the precise mechanism by which the PIDDosome forms specifically after mitosis and which cell cycle regulators control this process remain unclear. Understanding this mechanism is particularly important in the context of therapeutic strategies, as some cancer cells evade treatment by escaping cell cycle arrest. Although earlier studies have suggested that sensitising these cells to mitotic catastrophe could be a potential approach, the exact mechanism underlying this process remains poorly understood (Denisenko *et al*, 2016; Sazonova *et al*, 2021).

Here, we propose a probable mechanism of mitotic catastrophe based on experimentally validated biological interactions (**Fig. 6A**). For simplicity, we divide the proposed mitotic catastrophe module into two submodules: the first one is p53-induced activation of PIDD, and the second is Casp2 regulation by cell cycle regulators (**Fig. 6A**). In the first submodule, p53 activates its downstream target, PIDD. However, experimental studies suggest that PIDD activation occurs during DNA synthesis (S-phase) (Burigotto *et al*, 2021; Bock *et al*, 2012). To incorporate this interaction into our model phenomenologically, we assume that PIDD activation of PIDDact is inhibited by Cdt1, a protein expressed during the early phase of the cell cycle and responsible for initiating DNA replication (Ratnayeke *et al*, 2023) (**Fig. 6A**). Cdt1 is degraded as DNA synthesis begins, ensuring that active PIDD appears only after DNA synthesis begins. The second module focuses on Casp2, another component of the PIDDosome complex. Casp2 is present under normal conditions but remains inactive during the S-G_2_-M phase due to phosphorylation by AURKB and CDK1-CycB (Lim *et al*, 2021).

**Fig. 6.**
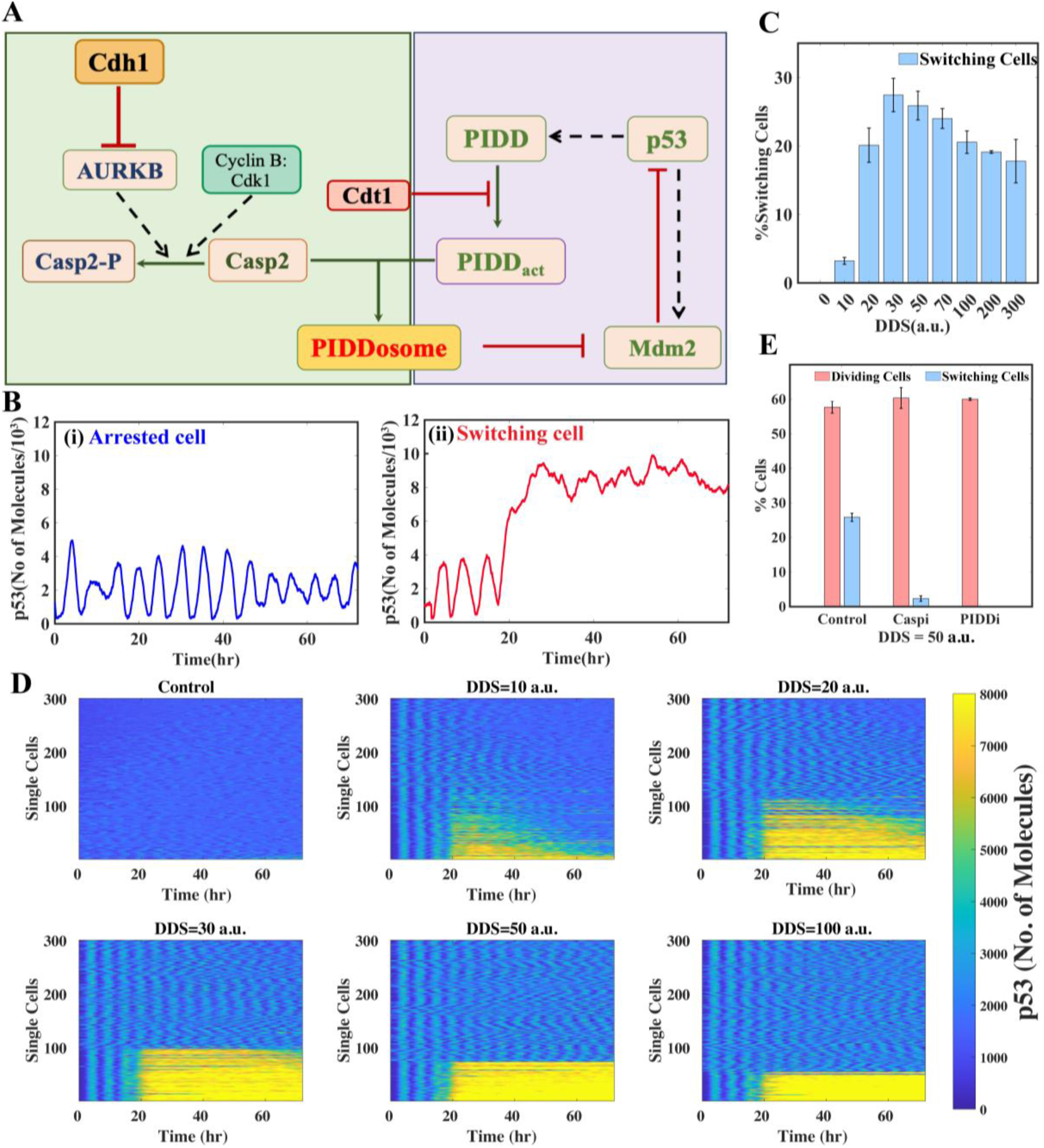
Proposed mitotic catastrophe module and post-mitotic switching in p53 dynamics. ***(A)*** Proposed mitotic catastrophe mechanism incorporating key p53 and cell cycle regulators. ***(B)*** Representative p53 time profile for ***(i)*** arrested cell and ***(ii)*** switching cell. ***(C)*** Percentage of cells exhibiting switching p53 dynamics at different DNA damage doses. ***(D)*** p53 expression profile at different DNA damage signals. (n=300 for each condition). ***(E)*** Fraction of switching and dividing cells under control condition, caspase inhibition, and PIDD inhibition at *DDS* = 50 a.u..

That means Casp2 will be suppressed during the S-G_2_-M phase of the cell cycle and will be present only to form a complex with PIDDact during the G_1_ phase (**Fig. 6A**). Based on these interactions, we hypothesise the following dynamics (**Fig. S3**). When a cell escapes G_1_ cell cycle arrest and enters S phase, p53-induced PIDD is activated during DNA synthesis (Burigotto *et al*, 2021; Bock *et al*, 2012). However, during the S-G_2_-M phase, because CDK1 remains active, Casp-2 remains inactive. After mitosis, CycB is degraded, and Casp2 is rapidly dephosphorylated and released. Active Casp2 can now bind to PIDDact to form the PIDDosome complex. The PIDDosome promotes degradation of Mdm2 (Oliver *et al*, 2011), leading to p53 stabilisation and triggering a sharp switch in p53 dynamics (**Fig. S3**). This mechanism provides a dynamical explanation for why the p53 switch occurs specifically after mitosis in the escaped cells.

### Model captures the post-mitosis switching in p53 dynamics

We have incorporated all additional interactions discussed above into our coupled network (**Fig. 1**) and simulated it within a stochastic framework. Our results demonstrate that cells escaping G_1_ arrest exhibit switch-like dynamics in p53 expression after mitosis (**Fig. 6B)**. In contrast, non-dividing cells show oscillatory dynamics of p53 (**Fig. 6B)**. This difference is evident in the average profiles as well, where escaper cells show clear p53 switching dynamics and non-dividing cells exhibit a damped oscillatory p53 time profile due to averaging of p53 oscillations (**Fig. S4)**. Interestingly, the fraction of switching cells shows a dose-dependent response to DNA damage. As the DNA damage dose increases, the number of switching cells rises, but declines after a certain threshold. To better understand this trend, we systematically investigated the dose dependence of p53-mediated mitotic catastrophe and the number of cell divisions (**Fig. 6C**). At low DNA-damage doses, cell divisions are more frequent. However, because there is less DNA damage, the repair process becomes faster (**Fig. S5A**). Consequently, the normal cell cycle will be restored, and the probability of a cell undergoing mitotic catastrophe will be lower. As the damage dose increases, repair takes longer (**Fig. S5A**). This prolonged exposure increases the likelihood of stochastic errors, thereby increasing the risk of mitotic catastrophe (**Fig. 6D**). However, at very high levels of DNA damage, p53 expression becomes strongly elevated (**Fig. S5B**). High p53 levels strongly enforce robust cell cycle arrest and reduce the number of cells undergoing mitotic catastrophe (**Fig. 6D**). To further investigate whether mitotic catastrophe depends on PIDD and caspase-2, we performed an inhibition simulation by reducing their levels. In both cases, the number of cells exhibiting mitotic catastrophe decreased significantly, without affecting the total number of dividing cells (**Fig. 6E**). These results support the idea that PIDD and Casp2 are both required to initiate mitotic catastrophe, which aligns with the experimental findings (Tsabar *et al*, 2020).

### Model predicts the ways to induce mitotic catastrophe to attain therapeutic advantage

The dose-response analysis (**Fig. 6C**) shows that the highest fraction of cells showing mitotic catastrophe is at the intermediate DNA damage level. At low doses, very few cells experience mitotic catastrophe, mainly because DNA damage repair is rapid (**Fig. S5A**), and most cells repair the damage before defective mitosis occurs. However, from a therapeutic perspective, it would be beneficial to induce mitotic catastrophe even at low levels of DNA damage, thereby inducing apoptosis in these cells. Based on our hypothesis, prolonging DNA repair time could increase the likelihood of mitotic catastrophe. To check this, we simulated the effect of a DNA damage repair inhibitor (DDRi) (Drew *et al*, 2025). Our simulations show that across all DNA-damage doses, the fraction of cells exhibiting mitotic catastrophe increases after DDRi compared with control (**Fig. 7A**). This suggests that DDRi, together with DNA damage, can be utilised to selectively eliminate cells that escape arrest, even at low DNA-damage doses.

**Fig. 7.**
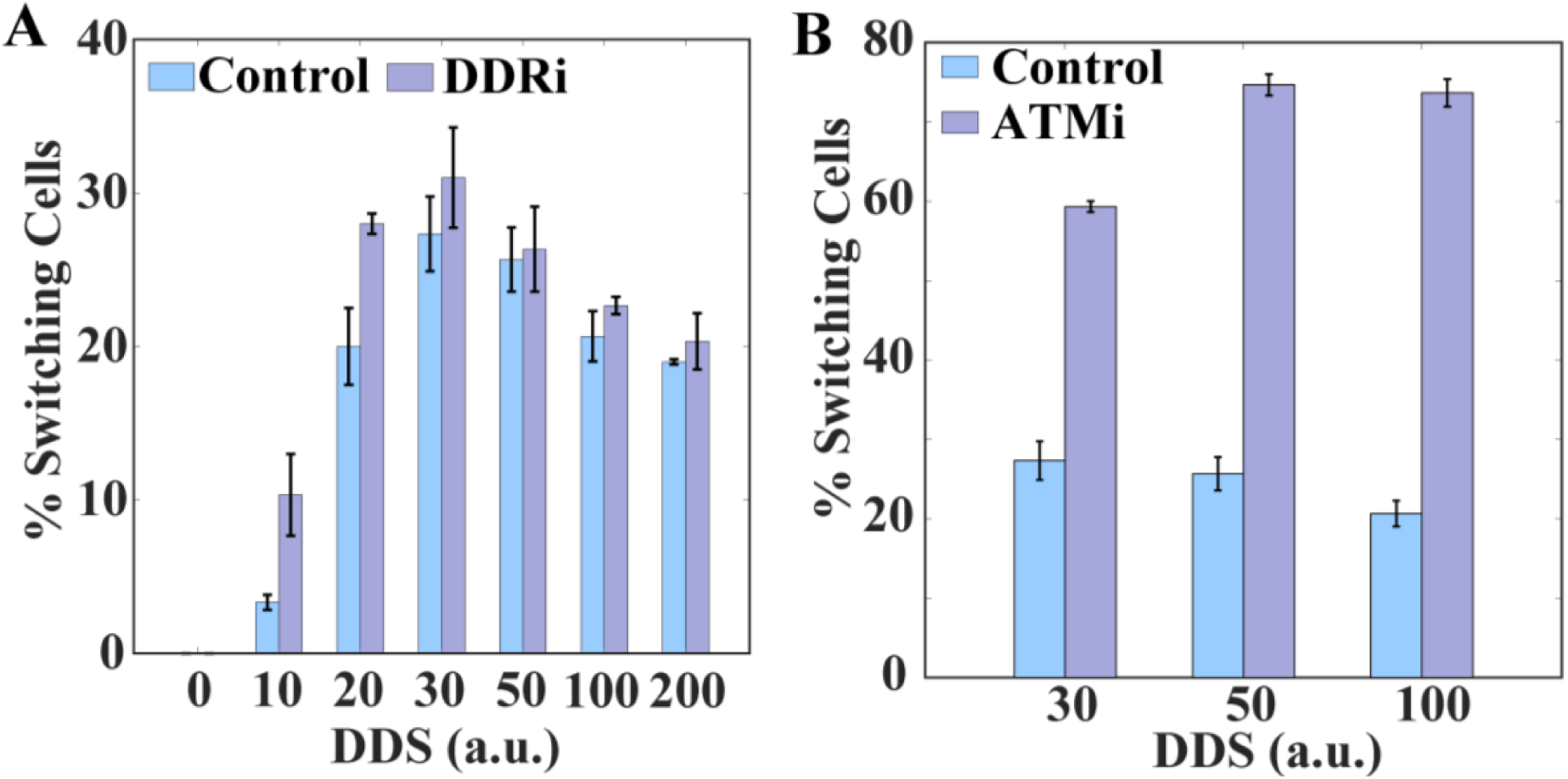
Model prediction for inducing mitotic catastrophe selectively. Percentage of switching cells while combining DNA damage signal with **(A)** DNA damage repair inhibitor (DDRi) and **(B)** ATM inhibitor (ATMi).

Next, we explored whether increasing the mitotic catastrophe could be enhanced at a fixed DNA damage dose and repair rate. One approach is to weaken the signal of DNA damage to p53. A slightly reduced p53 induces more cells to slip past cell cycle arrest and enter the cell cycle, where they could undergo mitotic catastrophe. To investigate this, we consider the ATM kinase, which is activated in response to DNA damage (Batchelor *et al*, 2011). ATM phosphorylates Mdm2, leading to its degradation (Cheng *et al*, 2011). In our model, this interaction is phenomenologically represented as DNA damage degrading Mdm2 (Charan *et al*, 2022). We simulated ATM inhibition (ATMi) by weakening this interaction. Our simulations reveal that ATMi increases the number of cells that escape G_1_ arrest due to low p53 levels. However, once these cells progress through the cell cycle and divide, mitotic catastrophe occurs (**Fig. 7B**). Importantly, ATM and PIDDosome regulate Mdm2 through distinct mechanisms: ATM degrades Mdm2 via phosphorylation at multiple sites, whereas the PIDDosome cleaves Mdm2 at Asp 367, leading to post-mitotic p53 stabilisation (Cheng *et al*, 2011; Oliver *et al*, 2011). Thus, even when ATM is reduced, the PIDDosome pathway can still trigger a sharp increase in p53. In summary, our model predicts that combinations of DNA damage with repair and checkpoint modulation by specific inhibitors could selectively induce mitotic catastrophe, thereby conferring a therapeutic advantage.

## Conclusion

p53 plays a central role in mediating the DNA damage signal by halting cell cycle progression to maintain genomic integrity (Levine, 1997; Vousden & Prives, 2009; Kastan *et al*, 1991). However, recent experimental studies show that p53 sometimes fails to maintain cell cycle arrest when the DNA damage signal persists for an extended period (Tran *et al*, 2023; Tsabar *et al*, 2020; Reyes *et al*, 2018). Under these circumstances, cells escape G_1_ arrest and proceed through mitosis despite having damaged DNA, resulting in sustained, high post-mitotic p53 levels and mitotic catastrophe (Tsabar *et al*, 2020; Tran *et al*, 2023). However, the key molecular regulator governing escape from the G1-arrested state and subsequent mitotic catastrophe remains unclear and has not been explored using mathematical modelling. In this article, we developed an integrated model that couples the core cell cycle regulatory network with p53 dynamics (**Fig.1**). Our model captures cell cycle arrest in response to DNA damage (**Fig. 2**). The stochastic simulations of it confirm that a subset of cells indeed exhibits escape from cell cycle arrest (**Fig. 3** and **Fig. 4A**) and the extent of escaping cells varies in DNA damage dose-dependent manner. Herein, we investigated which key molecular interactions underlie this escape phenomenon.

Through systematic analysis, our model predicts that escaper cells exhibit high levels of E2F1 (**Fig. 5B**). The presence of stochastic fluctuations often facilitates sudden CycE activation and allows such cells to escape the G_1_-S checkpoint despite having damaged DNA (**Fig. 5D**). This reflects the intricate interplay between cell cycle regulation and p53 signalling that determines cell fate decision-making. Our model further captures instances in which, among the escaping cells, some exhibit random geminin peaks during prolonged arrest (**Fig. 4A**) but do not divide. In these cells, sustained accumulation of E2F1 partially activates Cyclin A (**Fig.5B-C**). Partial Cyclin A activation leads to incomplete inactivation of Cdh1, allowing geminin to rise, without activating CycB (**Fig. 5B-C**). As a result, mitosis does not occur. This model finding correlates with several experimental studies, in which similar failed division attempts and endocycle-like events were observed under prolonged DNA damage conditions (Fujimaki *et al*, 2025; Tran *et al*, 2023). Thus, the model simulations elucidate the roles of key cell cycle regulators in controlling the fluctuating dynamics of escaper cells during DNA damage.

To explain post-mitotic p53 switching and its relation to mitotic catastrophe, we propose an additional regulatory module that further strengthens the cross-talk between cell cycle regulation and p53 dynamics (**Fig. 6A**). Consistent with Tsabar et al.’s finding, our model shows that when cells divide in the presence of DNA damage, the PIDDosome complex forms(Tsabar *et al*, 2020); this complex stabilises p53 by degrading Mdm2. The timing of PIDDosome formation ensures that mitotic catastrophe only occurs after mitotic exit in cells that have crossed the G_1_-S checkpoint in the presence of DNA damage (**Fig. S3**). This intricate and tight regulation signifies how robustly the cellular system functions. The model demonstrates that mitotic catastrophe exhibits DNA damage dose dependence, with intermediate doses leading to a higher fraction of cells exhibiting switching of p53 dynamics (**Fig. 6C**). It further illustrates that under low DNA-damage conditions, the faster repair has a very narrow time window for cells to escape (**Fig. 6D**). On the other hand, at higher DNA-damage doses, higher levels of p53 keep a tight hold on p21, preventing cells from facilitating escape. Based on these insights, our model predicts that combining DNA damage with a DNA repair inhibitor increases the fraction of cells undergoing mitotic catastrophe by prolonging DNA damage. Additionally, weakening ATM-mediated p53 activation allows more cells to escape arrest, which increases post-mitotic p53 activation through the PIDDosome pathway **(Fig. 7)**.

In summary, we propose an integrated cell-cycle and p53 regulatory network in response to DNA damage. Our results reveal that stochastic fluctuations and intrinsic levels of cell cycle regulators drive the escape from cell cycle arrest during prolonged DNA damage. The proposed mitotic catastrophe mechanism captures the intricate crosstalk between the cell cycle and p53 dynamics by modelling p53’s switching behaviour. Overall, our model provides a mechanistic understanding of cell cycle regulation in the presence of DNA damage and makes experimentally testable predictions with therapeutic implications.

### Limitations of the study

One limitation of our model is that the fraction of cells committing to the cell cycle within the first 24 hrs of DNA damage is high. This might be due to the simplified network, which excludes other checkpoints. To minimise the impact of that, we excluded cells that divided within 48 hours of DNA damage by analysing only those that divided after 48 hours of DNA damage.

## Method

### Deterministic simulations

The coupled network was described by ordinary differential equations (**Tables S1, S5, and S9**). Deterministic simulations of the model equations were performed using the freely available software XPPAUT (https://sites.pitt.edu/ phase/bard/bardware/xpp/xpp.html). In the deterministic simulations, all the variables were in scaled units (s.u.).

### Stochastic simulation

We have translated the deterministic mathematical model into a stochastic framework by considering the reactions shown in **Tables S4 & Table S8,** and simulated the system using Gillespie’s stochastic simulation algorithm (SSA)(Gillespie, 1976). The simulation was initiated with the same initial conditions 20 hours before DNA damage induction to make cells asynchronous before the DNA damage signal was applied.

### Classification of cells in single divider, non-divider, and Geminin random and protein expression calculation

Based on the mitosis profile and Geminin expression shown in **Fig. 4A**, cells were classified into three categories **Fig. 5A**. The first category consists of non-dividing cells, where there is no division and no high Geminin expression (threshold value = 3500 molecules). The second category includes the cells that escape the cell cycle arrest after 48 hours of DNA damage. The third category includes cells with high Geminin expression that do not undergo mitosis.

### p53 switching threshold calculation for mitotic catastrophe

To classify the cells as p53 switcher cells, we defined a threshold of 8000 molecules, which is approximately double the average of oscillatory p53. In addition, a temporal criterion was applied: the cell should maintain p53 levels above 7500 molecules for at least 20 hr. Cells that satisfied both the amplitude and duration thresholds were considered switching cells.

## Acknowledgments

Thanks are due to UGC for providing the UGC-CSIR-JRF (NTA Ref. No: 191620004555) fellowship to KC. This work is supported by the funding agency **ANRF (Erstwhile SERB), India** (Grant no. **CRG/2023/002165**).

## Notes

### Competing Interest Statement

The authors have declared no competing interest.

