## Supplementary material - Text, Figures and tables for "Intricate Dynamical Cross-Talk Between p53 Protein and Cell Cycle Regulators Governs Mammalian Cell Fate"

#### Supplementary Text

1. **Model description for mitotic catastrophe module -** The coupled model of p53 and cell cycle regulation is built by adapting previously established models of p53 and cell cycle regulatory networks (Charan *et al*, 2022; Govindaraj *et al*, 2022). However, the kinetic equations of the proposed mitotic catastrophe module are discussed in the following sections.

**Eqs. 1 and 2** describe the dynamics of PIDD (p53-induced death domain) protein and its activated form, respectively. The first term in **Eq. 1** describes the basal synthesis rate, whereas the second and third terms describe the p53 dimer and tetramer-mediated synthesis rate of PIDD protein. The third term includes the degradation of PIDD protein. The last term corresponds to PIDD activation, which occurs only during the DNA synthesis phase. We have phenomenologically introduced this effect by inhibiting this process by Cdt1.To ensure its activation during DNA damage, we have also included the DNA damage effect in the last term.

$$\frac{dPIDD}{dt} = k_{spi} + k_{pid} \times p53D + k_{pit} \times p53T - k_{dpi} \times PIDD - k_{pact} \times \frac{PIDD}{k_{gem} + PIDD} \times \left(\frac{1}{1 + kcdtc \times Cdt1}\right) \times \frac{DNA_{dam}}{1 + DNA_{dam}} \quad (\boldsymbol{Eq.1})$$

The first term in **Eq. 2** shows the activation into PIDDact form from the PIDD protein. The second term describes the degradation of the PIDDact form. The third and fourth terms correspond to the formation and dissociation of the PIDDosome (PDCa) heterodimer, respectively.

$$\boldsymbol{\frac{dPIDDact}{dt} = (k_{pact}) \times \frac{PIDD}{k_{gem} + PIDD} \times \left(\frac{1}{1 + kcdtc \times Cdt1}\right) \times \frac{DNA_{dam}}{1 + DNA_{dam}} - k_{dap} \times PIDD_{act} - k_{com} \times \frac{PIDDact}{k_{pdd} + PIDDact} \times \frac{casp2}{kpcc + casp2} + k_{depc} \times PDCa} \quad (\boldsymbol{Eq.2})$$

**Eqs. 3 and 4** show the dynamics of the Caspase-2 protein in its active and phosphorylated forms. The first term in **Eq. 3** denotes the synthesis of casp-2. The subsequent four terms show the phosphorylation of Casp-2 into Casp2i by basal, CycA, CycB, and AURKB phosphorylation. The fifth term describes the dephosphorylation of Casp-2. The sixth term shows the degradation of Casp-2 in non-phosphorylated form. The last two terms describe the formation and dissociation of PDca complex from PIDDact and Casp2, respectively.

$$\frac{dcasp2}{dt} = k_{scasp} - \frac{k_{11s} \times casp2}{J_{11} + +casp2} - \frac{k_{11a} \times casp2 \times CycA}{J_{12} + casp2} - \frac{k_{11b}\, casp2 \times CycBa}{J_{13} + casp2} - \frac{k_{11c} \times casp2 \times AURKB}{J_{14} + +casp2} + \frac{k_{11d} \times (casp2i)}{J_{15} + (casp2i))} - k_{dcasp} \times casp2 - k_{com} \times \frac{PIDDact}{k_{pdd} + PIDDact} \times \frac{casp2}{kpcc + casp2} + k_{depc} \times PDCa \qquad (Eq.3)$$

**In Eq. 4**, the first four terms describe the phosphorylation of Casp-2 into Casp2i. The fifth term denotes the dephosphorylation of Casp2i. The last term represents the degradation rate of Casp2i in its phosphorylated form.

$$\frac{dcasp2i}{dt} = \frac{k_{11s} \times casp2}{J_{11} + +casp2} + \frac{k_{11a} \times casp2 \times CycA}{J_{12} + casp2} + \frac{k_{11b}\, casp2 \times CycBa}{J_{13} + casp2} + \frac{k_{11c} \times casp2 \times AURKB}{J_{14} + +casp2} - \frac{k_{11d} \times (casp2i)}{J_{15} + (casp2i))} - k_{dcasp} \times casp2i \qquad (Eq.4)$$

**Eq. 5** describes the dynamics of the PIDDosome complex. The first term shows the formation of a complex from Casp2 and PIDDact. The second term denotes the dissociation of the PIDDosome complex. The last term represents the degradation of the PIDDosome complex.

$$\frac{dPDCa}{dt} = k_{com} \times \frac{PIDDact}{k_{pdd} + PIDDact} \times \frac{casp2}{kpcc + casp2} - k_{depc} \times PDCa - k_{degpc} \times \frac{PDca}{j_{deg} + PDca} \qquad (Eq.5)$$

**Eq. 6** describes the dynamics of the AURKB protein. The first term shows the synthesis of AURKB protein. The second and third terms represent the degradation of the AURKB protein, independent of and dependent on Cdh1, respectively.

$$\frac{dAURKB}{dt} = ksau - kdau \times AURK - kdauc \times AURKB \times \frac{Cdh1}{jau + AURKB} \qquad (Eq.6)$$

**Eq. 7** is the modified equation of Mdm2 after incorporating the PIDDosome-mediated degradation of Mdm2 in the last term.

$$\frac{dMdm}{dt} = k_{smdm} + k_{mono} \times P53_{mono} + k_{dimer} \times p53_2 + k_{tetramer} \times p53_4 + k_2 \times Mdm^p - (k_{dmdm} + \frac{k_{dnam} \times DNAdam}{k_a + DNAdam}) \times Mdm - k_1 \times Mdm - k_{PIDDosome} \times Mdm \times \frac{PDca^{mn}}{kpidd^{mn} + PDca^{mn}} \qquad (Eq.7)$$

2. **p53 and p21 siRNA simulations -** To inhibit p53 and p21, similar to siRNA, we modified the dynamical equations for both proteins in our simulations.

**In Eq. 8**, in the first term, the basal transcription rate is reduced by the p53si term in the denominator, which represents the reduced transcription.

$$\frac{dp53m}{dt} = \frac{J_0}{1+p53si} - k_{dpm} \times p53m \qquad (\boldsymbol{Eq.8})$$

Similarly, in **Eq. 9**, the first three terms representing the transcription of p21 were reduced by the p21si term in the denominator.

$$\frac{dI_m}{dt} = \frac{ksim}{1+p21si} + \frac{kp53d}{1+p21si} \times p53d + \frac{kp53t}{1+p21si} \times p53t - k_{dim} \times I_m \qquad (\boldsymbol{Eq.9})$$

**PIDD and casp inhibition** – The inhibition of PIDD and Casp2 were achieved by respectively modifying their dynamical equations.

The PIDD inhibitor is introduced in the denominator of PIDD activation (**Eq. 10**).

$$\frac{dPIDDact}{dt} = \frac{kpact}{1+PIDDi} \times \frac{PIDD}{k_{gem}+PIDD} \times \left(\frac{1}{1+kcdtc \times Cdt1}\right) \times \frac{DNA_{dam}}{1+DNA_{dam}} - k_{dap} \times PIDD_{act} - k_{com} \times \frac{PIDDact}{k_{pdd}+PIDDact} \times \frac{casp2}{kpcc+casp2} + k_{depc} \times PDCa \qquad (\boldsymbol{Eq.10})$$

The Casp2 inhibition is achieved by inhibiting the synthesis term of Caspase by introducing the inhibition term in the denominator (**Eq. 11**).

$$\frac{dcasp2}{dt} = \frac{k_{scasp}}{1+Caspi} - \frac{k_{11s} \times casp2}{J_{11}++casp2} - \frac{k_{11a} \times casp2 \times CycA}{J_{12}+casp2} - \frac{k_{11b}\, casp2 \times CycBa}{J_{13}+casp2} - \frac{k_{11c} \times casp2 \times AURKB}{J_{14}++casp2} + \frac{k_{11d} \times (casp2i)}{J_{15}+(casp2i))} - k_{dcasp} \times casp2 - k_{com} \times \frac{PIDDact}{k_{pdd}+PIDDact} \times casp2 + k_{depc} \times PDCa \qquad (\boldsymbol{Eq.11})$$

#### Supplementary Figures

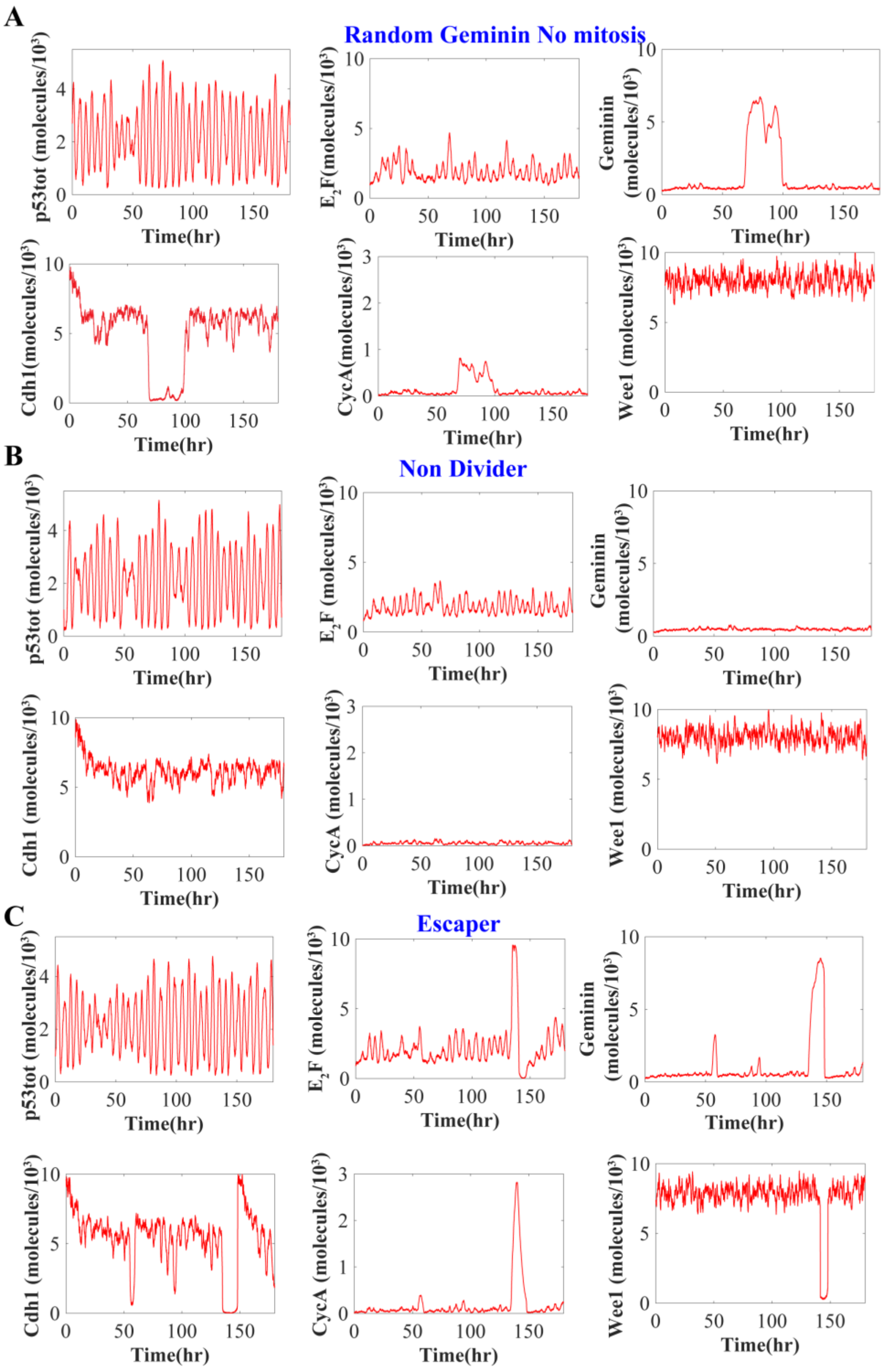

**Fig S1** – Representative time profiles of p53 total, E2F, Geminin, Cdh1, CycA, and Wee1 for a cell showing (A) Random geminin peak without mitosis, (B) No division, and (C) Escape from cell cycle arrest.

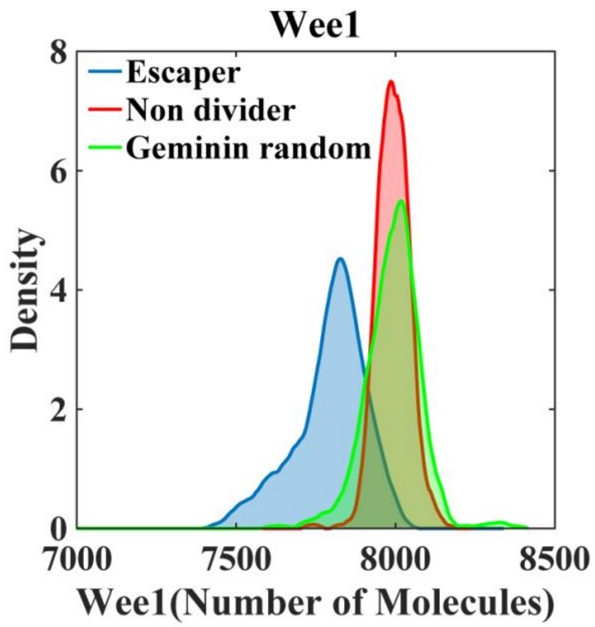

**Fig. S2** Expression profile of Wee1 in three cell categories.

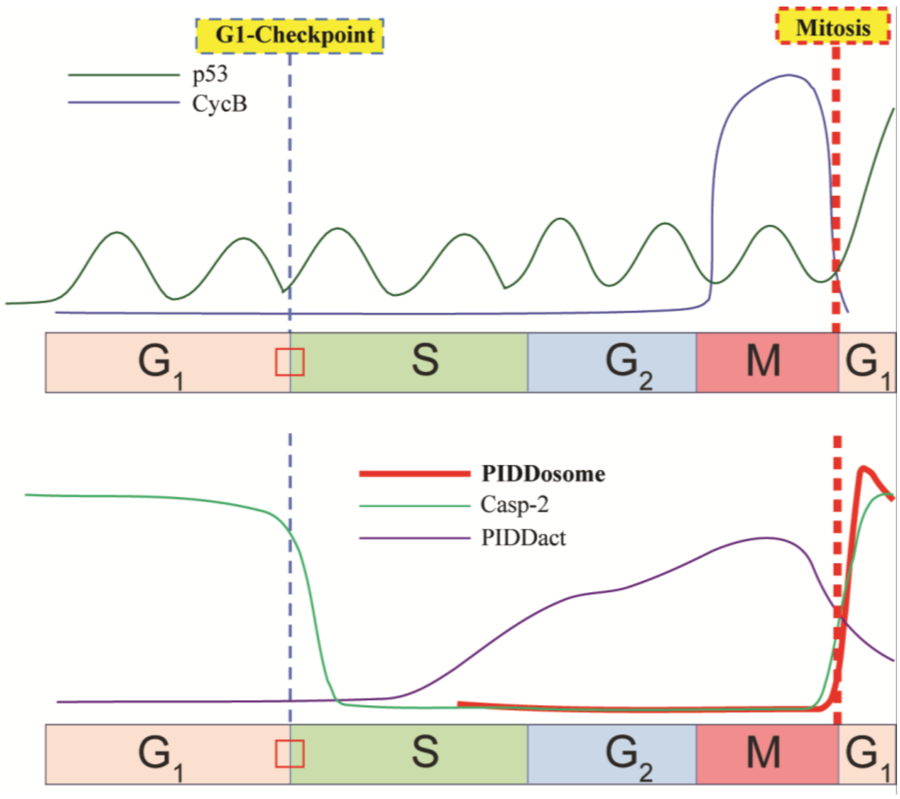

**Fig. S3** Proposed dynamics of p53 and CycB showing the $G_1$-S checkpoint escape and mitosis (top) and PIDDosome, Casp-2, and PIDDact during cell cycle progression (Bottom).

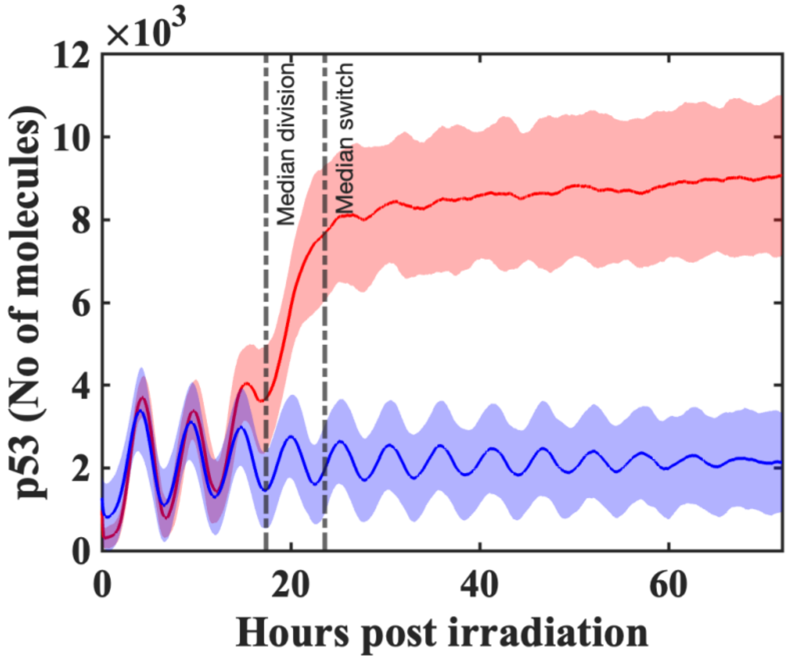

**Fig. S4** p53 dynamics in p53 switcher (red) and non-switcher (blue) cells at DDS=50. The solid line indicates the moving mean of p53, and the cloud represents the standard deviation.

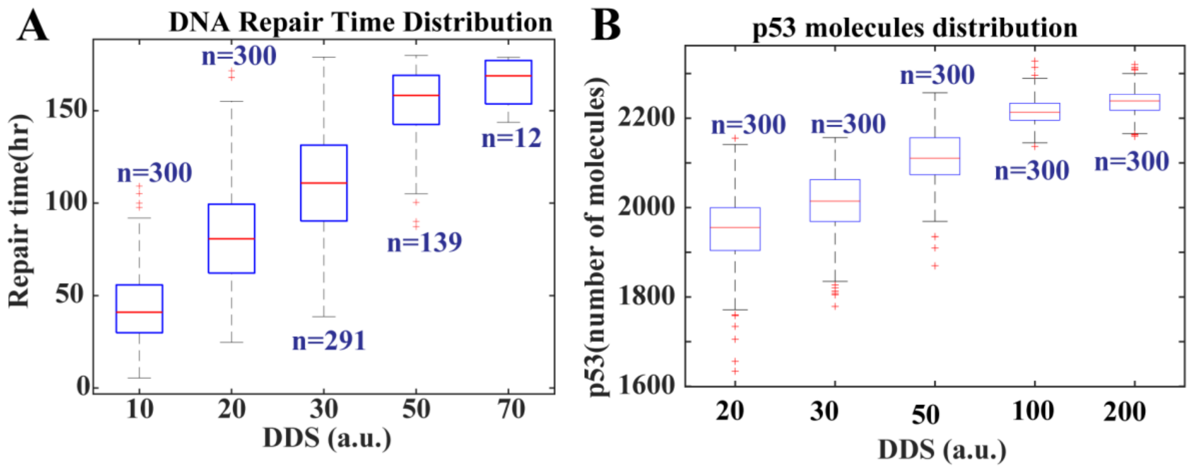

**Fig. S5 (A)** DNA damage repair time at different DNA damage signal dosages. **(B)** p53 mean level at different DNA damage signal dosages (n represents the number of cells).

### Supplementary Tables

**Table S1:** ODEs governing the dynamics of the p53 regulatory network. **(Charan K. et al., 2023 ChemPhysChem)**

| | | |
|---|---|---|
| $\frac{dp53m}{dt}$ | $= J_0 - k_{dpm} \times p53m$ | 1 |
| $\frac{dp53_{tot}}{dt}$ | $= k_{synp} \times p53m - k_{dm} \times p53_{mono} - \frac{k_{dyn} \times p53_{mono} \times Mdm^p}{k_{mm} + p53_{mono}} - 2 \times k_{dd} \times p53_2 - \frac{2 \times k_{dyn} \times p53_2 \times Mdm^p}{k_{mm1} + p53_2} - \frac{4 \times k_{dyn} \times p53_4 \times Mdm^p}{k_{mm2} + p53_4} - 4 \times k_{dt} \times p53_4$ | 2 |
| $\frac{dp53_2}{dt}$ | $= k_c \times p53_{mono}^2 - k_d \times p53_2 - (k_e + ktdna \times DNAdam) \times p53_2^2 + k_f \times p53_4 - k_{dd} \times p53_2 - \frac{k_{dyn} \times p53_2 \times Mdm^p}{k_{mm1} + p53_2}$ | 3 |
| $\frac{dp53_4}{dt}$ | $= (k_e + ktdna \times DNAdam) \times p53_2^2 - k_f \times p53_4 - \frac{k_{dyn} \times p53_4 \times Mdm^p}{k_{mm2} + p53_4} - k_{dt} \times p53_4$ | 4 |
| $\frac{dMdm}{dt}$ | $= k_{smdm} + k_{mono} \times P53_{mono} + k_{dimer} \times p53_2 + k_{tetramer} \times p53_4 + k_2 \times Mdm^p - (k_{dmdm} + \frac{k_{dnam} \times DNAdam}{k_a + DNAdam}) \times Mdm - k_1 \times Mdm$ | 5 |
| $\frac{dMdm^P}{dt}$ | $= k_1 \times Mdm - k_2 \times Mdm^p$ | 6 |
| $\frac{dDNAdam}{dt}$ | $= k_{int} + k_{dna} \times DDS - k_{ddna} \times \frac{DNAdam}{J_{dna} + DNAdam}$ | 7 |
| $p53_{mono}$ | $= p53_{tot} - (2 \times p53_2 + 4 \times p53_4)$ | 8 |

Note: all the equations in Table-S1 are multiplied by scaling factor 60 to change the timescale from minutes to hours.

**Table S2:** Abbreviated names and description of the variables in the p53 regulatory network. **(Charan K. et al., 2023 ChemPhysChem)**

| Abbreviated names | Description |
|---|---|
| $\boldsymbol{p53m}$ | $p53$ mRNA |
| $\boldsymbol{p53_{mono}}$ | $p53$ monomer |
| $\boldsymbol{p53_2}$ | $p53$ dimer |
| $\boldsymbol{p53_4}$ | $p53$ tetramer |
| $\boldsymbol{Mdm}$ | $Mdm2$ protein in unphosphorylated form |
| $\boldsymbol{Mdm^p}$ | $Mdm2$ protein in the phosphorylated form |
| $\boldsymbol{DNAdam}$ | DNA damage foci |

**Table S3**: Description and values of the parameters used in the p53 regulatory network. **(Charan K. et al., 2023 ChemPhysChem)**

| Symbol | Description | Value and unit |
|---|---|---|
| $\boldsymbol{k_{dyn}}$ | $Mdm^p$ dependent p53 degradation | $0.375\ min^{-1}$ |
| $\boldsymbol{J_0}$ | The basal synthesis rate of p53mRNA | $0.25\ \text{s.u.}min^{-1}$ |
| $\boldsymbol{k_{dpm}}$ | The degradation rate of p53mRNA | $0.03\ min^{-1}$ |
| $\boldsymbol{k_{synp}}$ | The synthesis rate of p53 | $1.25\ min^{-1}$ |
| $\boldsymbol{k_{mm}}$ | Michalis-Menten constant for degradation of $p53_{mono}$ | $1.0\ \text{s.u}$ |
| $\boldsymbol{k_{mm1}}$ | Michalis-Menten constant for degradation of $p53_2$ | $30.0\ \text{s.u}$ |
| $\boldsymbol{k_{mm2}}$ | Michalis-Menten constant for degradation of $p53_4$ | $1.0\ \text{s.u}$ |
| $\boldsymbol{k_{dm}}$ | The degradation rate of $p53_{mono}$ | $0.0002\ min^{-1}$ |
| $\boldsymbol{k_{dd}}$ | The degradation rate of $p53_2$ | $0.00025\ min^{-1}$ |
| $\boldsymbol{k_{dt}}$ | The degradation rate of $p53_4$ | $0.002\ min^{-1}$ |
| $\boldsymbol{k_c}$ | On rate for Dimerization reaction | $0.012\ \text{s.u}^{-1}.min^{-1}$ |
| $\boldsymbol{k_d}$ | The off rate for Dimerization reaction | $0.0000013\ min^{-1}$ |
| $\boldsymbol{k_e}$ | On rate for tetramerization reaction | $0.0002\ \text{s.u}^{-1}.min^{-1}$ |

| | | |
|---|---|---|
| $k_f$ | The off rate for tetramerization reaction | $10^{-6}\ min^{-1}$ |
| $k_{smdm}$ | Basal synthesis rate for Mdm2 | $0.0025\ \mathrm{s.u.}min^{-1}$ |
| $k_{tetramer}$ | $P53_4$ activated mdm2 synthesis rate | $0.016\ min^{-1}$ |
| $k_{dmdm}$ | Mdm2 degradation rate | $0.000\ min^{-1}$ |
| $k_1$ | Phosphorylation rate of Mdm2 | $0.0017\ min^{-1}$ |
| $k_2$ | Dephosphorylation rate of $Mdm2^P$ | $0.017\ min^{-1}$ |
| $k_{dnam}$ | DNA damage mediated Mdm2 degradation rate | $0.015\ min^{-1}$ |
| $k_{dna}$ | DDS activated synthesis rate of DNA damage | $0.5\ min^{-1}$ |
| $k_{ddna}$ | DNA damage repair rate | $0.005\ \mathrm{s.u}^{-1}.min^{-1}$ |
| $k_{tdna}$ | DNA damage activated tetramerization rate | $0.0025\mathrm{s.u}^{-1}.min^{-1}$ |
| $DDS$ | DNA damage signal | 0 - 300 a.u. |
| $J_{dna}$ | Michaelis Menten constant in DNA damage repair | 1.0 |
| $k_a$ | Saturation term for Mdm2 degradation by DNA damage | 1.5 |
| $k_{mono}$ | P53 Monomer activated Mdm2 synthesis | $0.000007\ min^{-1}$ |
| $k_{dimer}$ | P53Dimer activated Mdm2 synthesis | $0.00005\ min^{-1}$ |
| $k_{int}$ | Intrinsic DNA damage | $0.0005\ min^{-1}$ |

**Table- S4**: Reactions governing the p53 regulatory network. **(Charan K. et al., 2023 ChemPhysChem)**

| Reaction number | Reaction | Propensity function |
|---|---|---|
| 1 | $\longrightarrow p53m$ | $w1 = J_0$ |
| 2 | $p53m \longrightarrow$ | $w2 = k_{dpm} \times p53m$ |
| 3 | $\xrightarrow{p53m} p53_{mono}$ | $w3 = k_{synp} \times p53m$ |

| | | |
|---|---|---|
| **4** | $p53_{mono} \longrightarrow$ | $w4 = k_{dm} \times p53_{mono}$ |
| **5** | $p53_{mono} + p53_{mono} \longrightarrow p53_2$ | $w5 = k_c \times p53_{mono} \times (p53_{mono} - 1)$ |
| **6** | $p53_2 \longrightarrow p53_{mono} + p53_{mono}$ | $w6 = k_d \times p53_2$ |
| **7** | $p53_{mono} \xrightarrow{Mdm^p}$ | $w7 = \frac{k_{dyn} \times p53_{mono} \times Mdm^p}{k_{mm} + p53_{mono}}$ |
| **8** | $p53_2 + p53_2 \longrightarrow p53_4$ | $w8 = k_e \times p53_2 \times (p53_2 - 1)$ |
| **9** | $p53_2 + p53_2 \xrightarrow{DNAdam} p53_4$ | $w9 = k_e \times p53_2 \times (p53_2 - 1) \times DNAdam$ |
| **10** | $p53_4 \longrightarrow p53_2 + p53_2$ | $w10 = k_f \times p53_4$ |
| **11** | $p53_2 \longrightarrow$ | $w11 = k_{dd} \times p53_2$ |
| **12** | $p53_2 \xrightarrow{Mdm^p}$ | $w12 = \frac{k_{dyn} \times p53_2 \times Mdm^p}{k_{mm1} + p53_2}$ |
| **13** | $p53_4 \longrightarrow$ | $w13 = k_{dt} \times p53_4$ |
| **14** | $p53_4 \xrightarrow{Mdm^p}$ | $w14 = \frac{k_{dyn} \times p53_4 \times Mdm^p}{k_{mm1} + p53_4}$ |
| **15** | $\longrightarrow Mdm$ | $w15 = k_{smdm}$ |
| **16** | $\xrightarrow{p53tot} Mdm$ | $w16 = k_{mono} \times p53_{mono} + k_{dimer} \times p53_2 + k_{tetramer} \times p53_4$ |
| **17** | $Mdm \longrightarrow$ | $w17 = k_{dmdm} \times Mdm$ |
| **18** | $Mdm \xrightarrow{DNAdam}$ | $w18 = \frac{k_{dnam} \times DNAdam \times Mdm}{k_a + DNAdam}$ |
| **19** | $Mdm \longrightarrow Mdm^p$ | $w19 = k_1 \times Mdm$ |
| **20** | $Mdm^p \longrightarrow Mdm$ | $w20 = k_2 \times Mdm^p$ |
| **21** | $\longrightarrow DNAdam$ | $w21 = k_{int}$ |
| **22** | $\xrightarrow{DDS} DNAdam$ | $w22 = k_{dna} \times DDS$ |
| **23** | $DNAdam \rightarrow$ | $w23 = \frac{k_{ddna} \times DNAdam}{J_{dna} + DNAdam}$ |

**Table - S5**. Equations governing reactions for the cell cycle network. **(Govindaraj et al., 2022 ACS Synth. Biol.)**

| | |
|---|---|
| $\frac{dMV}{dt} = \frac{k_{mV} \times s \times Serum}{cf \times (smv + (Serum * gmv))} - k_{dmv} \times MV$ | 1 |
| $\frac{dMyc_m}{dt} = (b_{myc} \times s) + k_{Mcm1} \times MV + \frac{k_{Mcm2} \times s \times DE^n}{(k_{mm}^n \times s^n) + DE^n} - k_{dMcm} \times Myc_m$ | 2 |
| $\frac{dMyc_p}{dt} = k_{Mcp} \times Myc_m - k_{dMcp} \times Myc_p - \frac{k_{dMcp1} \times Myc_p \times Skp2_a}{(s \times J_{ms}) + Myc_p}$ | 3 |
| $\frac{dE2F_m}{dt} = (b_{e2f} \times s) + k_{Em1} \times Myc_p + \frac{k_{Em2} \times s \times DE^n}{(k_{mm}^n \times s^n) + DE^n} - k_{dEm} \times E2F_m$ | 4 |
| $\frac{dE2F_T}{dt} = k_{Ep1} \times E2F_m - k_{dEp} \times E2F_p - k_{dEp} \times DE - k_{dEp} \times iDE - \frac{k_{dEp1} \times CycA \times E2F_p}{(s \times J_{DE}) + E2F_p} - \frac{k_{dEp1} \times CycA \times DE}{(s \times J_{DE}) + DE}$ | 5 |
| $\frac{dDE}{dt} = \frac{k_{DE1}}{s} \times Dp1_p \times E2F_p - k_{DE2} \times DE - \frac{k_{RD1}}{s} \times DE \times Rb + k_{RD2} \times iDE - k_{dEp} \times DE + \frac{k_{iR3} \times iDEP \times CycE}{(J_{RE} \times s) + iDEP} - \frac{k_{RpD1}}{s} \times DE \times iRb1 + k_{RpD2} \times iDEP - \frac{k_{dEp1} \times CycA \times DE}{(s \times J_{DE}) + DE}$ | 6 |
| $\frac{diDE}{dt} = \frac{k_{RD1}}{s} \times DE \times Rb - k_{RD2} \times iDE - k_{dEp} \times iDE - \frac{k_{iR1} \times iDE \times CycD}{(J_{RD} \times s) + iDE} + \frac{k_{iR2} \times s \times iDEP}{(J_{R1} \times s) + iDEP}$ | 7 |
| $\frac{diDE}{dt} = \frac{k_{RD1}}{s} \times DE \times Rb - k_{RD2} \times iDE - k_{dEp} \times iDE - \frac{k_{iR1} \times iDE \times CycD}{(J_{RD} \times s) + iDE} + \frac{k_{iR2} \times s \times iDEP}{(J_{R1} \times s) + iDEP}$ | 8 |
| $\frac{dRb_m}{dt} = k_{Rbm} \times s - k_{dRbm} \times Rb_m$ | 9 |
| $\frac{dRb_T}{dt} = k_{Rbp} \times Rb_m - k_{dRbp} \times (Rb + iRb1 + iRb2) - k_{dEp} \times iDE$ | 10 |
| $\frac{diRb1}{dt} = \frac{k_{iR1} \times Rb \times CycD}{(J_{RD} \times s) + Rb} - \frac{k_{iR2} \times s \times iRb1}{(J_{R1} \times s) + iRb1} - \frac{k_{iR3} \times iRb1 \times CycE}{(J_{RE} \times s) + iRb1} + \frac{k_{iR4} \times s \times iRb2}{(J_{R2} \times s) + iRb2} - \frac{k_{RpD1}}{s} \times DE \times iRb1 + k_{RpD2} \times iDEP - k_{dRbp} \times iRb1$ | 11 |
| $\frac{diRb2}{dt} = \frac{k_{iR3} \times iRb1 \times CycE}{(J_{RE} \times s) + iRb1} + \frac{k_{iR4} \times s \times iRb2}{(J_{R2} \times s) + iRb2} + \frac{k_{iR3} \times iDEP \times CycE}{(J_{RE} \times s) + iDEP} - k_{dRbp} \times iRb2$ | 12 |
| $\frac{dCycD_m}{dt} = k_{CDm1} \times MV + k_{CDm2} \times Myc_p - k_{dCDm} \times CycD_m$ | 13 |
| $\frac{dCycD}{dt} = k_{CDp1} \times CycD_m - k_{dCDp} \times CycD - \frac{k_{dCDp1} \times CycD \times CycA}{(s \times J_{CD}) + CycD}$ | 14 |
| $\frac{dCycE_m}{dt} = (b_{Cycem} \times s) + k_{sem} \times DE - k_{dCEm} \times CycE_m$ | 15 |
| $\frac{dCycE}{dt} = k_{se} \times CycE_m - k_{dE} \times CycE - \frac{k_{dEA}}{s} \times CycE \times Skp2 - \frac{k_{aei}}{s} \times CycE \times I + k_{dei} \times EI + \frac{k_{dip}}{s} \times EI \times Skp2_a$ | 16 |
| $\frac{dI_m}{dt} = ksim + kp53d \times p53d + kp53t \times p53t - k_{dim} \times I_m$ | 17 |
| $\frac{dI}{dt} = k_{si} \times I_m - k_{di} \times I - \frac{k_{ip} \times CycE \times I}{(J_{CE} \times s) + I} - \frac{k_{aei}}{s} \times CycE \times I + k_{dei} \times EI$ | 18 |
| $\frac{dI_p}{dt} = \frac{k_{ip} \times CycE \times I}{(J_{CE} \times s) + I} - k_{dip1} \times I_p - \frac{k_{dip}}{s} \times I_p \times Skp2_a$ | 19 |
| $\frac{dEI}{dt} = \frac{k_{aei}}{s} \times CycE \times I - k_{dei} \times EI - k_{di} \times EI - k_{dE} \times EI - \frac{k_{dip}}{s} \times EI \times Skp2_a$ | 20 |

| | |
|---|---|
| $\frac{dSkp2_m}{dt} = k_{17m} \times s - k_{17dm} \times Skp2_m$ | 21 |
| $\frac{dSkp2_T}{dt} = k_{17} \times Skp2_m - k_{18a} \times (Skp2_T - Skp2_a) - \frac{k_{18b} \times (Skp2_T - Skp2_a) \times Cdh1_a}{(J_{18} \times s) + Skp2_T - Skp2_a} - k_{18c} \times Skp2_a - \frac{k_{18d} \times Skp2_a \times Cdh1_a}{(J_{18b} \times s) + Skp2_a}$ | 22 |
| $\frac{dSkp2_a}{dt} = \frac{k_{19a}}{s} \times CycE \times (Skp2_T - Skp2_a) - k_{19b} \times Skp2_a - k_{18c} \times Skp2_a - \frac{k_{18d} \times Skp2_a \times Cdh1_a}{(J_{18b} \times s) + Skp2_a}$ | 23 |
| $\frac{dCycA_m}{dt} = b_{sam} \times s + k_{sam} \times DE - k_{dam} \times CycA_m$ | 24 |
| $\frac{dCycA}{dt} = k_{sa} \times CycA_m - k_{da} \times CycA - \frac{k_{da1}}{s} \times CycA \times Cdh1_a$ | 25 |
| $\frac{dCdh1_m}{dt} = k_{3m} \times s - k_{3dm} \times Cdh1_m$ | 26 |
| $\frac{dCdh1_T}{dt} = k_3 \times Cdh1_m - k_{3d} \times Cdh1_T$ | 27 |
| $\frac{dCdh1_a}{dt} = \frac{((k_{3a} \times s) + k_{3b} \times Cdc20_a) \times (Cdh1_T - Cdh1_a)}{(J_3 \times s) + Cdh1_T - Cdh1_a} - \frac{k_4 \times CycB_a \times Cdh1_a}{(J_4 \times s) + Cdh1_a} - \frac{k_{4b} \times CycA \times Cdh1_a}{(J_4 \times s) + Cdh1_a} - k_{3d} \times Cdh1_a$ | 28 |
| $\frac{dCdc20_m}{dt} = k_{5am} \times s + \frac{k_{5bm} \times s \times \left(\frac{CycB_a}{J_5 \times s}\right)^n}{1 + \left(\frac{CycB_a}{(J_5 \times s)}\right)^n} - k_{5dm} \times Cdc20_m$ | 29 |
| $\frac{dCdc20_T}{dt} = k_{5a} \times Cdc20_m - k_6 \times Cdc20_T$ | 30 |
| $\frac{dCdc20_a}{dt} = \frac{k_7 \times IEP \times (Cdc20_T - Cdc20_a)}{(J_7 \times s) + Cdc20_T - Cdc20_a} - \frac{k_8 \times Mad \times s \times Cdc20_a}{(J_8 \times s) + Cdc20_a} - k_6 \times Cdc20_a$ | 31 |
| $\frac{dIEP_a}{dt} = \frac{k_9}{s} \times CycB_a \times ((IEP_T \times s) - IEP_a) - k_{10} \times IEP_a$ | 32 |
| $\frac{dCycB_m}{dt} = k_{scbm} \times CycA - k_{dcbm} \times CycB_m$ | 33 |
| $\frac{dCycB_T}{dt} = k_{scb} \times CycB_m - k_{dcb} \times CycB_T - \frac{k_{dcb1}}{s} \times Cdh1_a \times CycB_T$ | 34 |
| $\frac{dCycB_a}{dt} = k_{scb} \times CycB_m + \frac{k_{cba}}{s} \times C25_a \times CycB_i - \frac{k_{cbi}}{s} \times Wee1_a \times CycB_a - k_{dcb} \times CycB_a - \frac{k_{dcb1}}{s} \times Cdh1_a \times CycB_a$ | 35 |
| $\frac{dC25_m}{dt} = k_{sc25m} \times CycA - k_{dc25m} \times C25_m$ | 36 |
| $\frac{dC25_T}{dt} = k_{sc25} \times C25_m - k_{dc25} \times C25_T$ | 37 |
| $\frac{dC25_a}{dt} = \frac{k_{c25a} \times C25_i \times CycB_a}{(k_{mc1} \times s) + C25_i} - \frac{k_{c25i} \times s \times C25_a}{(k_{mc2} \times s) + C25_a} - k_{dc25} \times C25_a$ | 38 |
| $\frac{dWee1_m}{dt} = \frac{k_{swm} \times s \times Serum}{sgw + (Serum * kgw)} - k_{dwm} \times Wee1_m$ | 39 |
| $\frac{dWee1_T}{dt} = k_{sw} \times Wee1_m - k_{dw} \times Wee1_T$ | 40 |
| $\frac{dWee1_a}{dt} = k_{sw} \times Wee1_m + \frac{k_{wa} \times s \times Wee1_i}{(k_{mc1} \times s) + Wee1_i} - \frac{k_{wi} \times Wee1_a \times CycB_a}{(k_{mw2} \times s) + Wee1_a} - k_{dw} \times Wee1_a$ | 41 |
| $\frac{dCdt1_m}{dt} = k_{21m} \times s - k_{21dm} \times Cdt1_m$ | 42 |

| | |
|---|---|
| $\frac{dCdt1}{dt} = k_{21} \times Cdt1_m - k_{22a} \times Cdt1 - \frac{k_{22b} \times Cdt1 \times Skp2_a}{(J_{22} \times s) + Cdt1}$ | 43 |
| $\frac{dGem_m}{dt} = k_{20m} \times s - k_{20dm} \times Gem_m$ | 44 |
| $\frac{dGem}{dt} = k_{19} \times Gem_m - k_{20a} \times Gem - \frac{k_{20b} \times Gem \times Cdh1_a}{(J_{20} \times s) + Gem}$ | 45 |
| Algebraic equations | |
| $E2F_p = E2F_T - DE - iDE - iDEP$; $Dp1_p = (Dp_t \times s) - DE - iDE - iDEP$; $Rb = Rb_T - iRb1 - iRb2 - iDE - iDEP$ | |
| $CycB_i = CycB_T - CycB_a$; $C25_i = C25_T - C25_a$; $Wee1_i = Wee1_T - Wee1_a$ | |

**Table- S6** Description of the model parameters and their values for cell cycle network **(Govindaraj et al., 2022 ACS Synth. Biol.)**

| Description | Parameter | Values | Unit |
|---|---|---|---|
| Serum | *Serum* | 2 | % |
| Synthesis rate of MV Protein | $k_{mv}$ | 3 | s.u. $h^{-1}$ |
| Michaelis-Menten constant associated with MV synthesis | *smv* | 2.2 | - |
| General serum mediated activation rate of MV synthesis | *gmv* | 0.2 | - |
| Degradation rate of MV protein | $k_{dmv}$ | 4 | $h^{-1}$ |
| Basal transcription rate of Myc mRNA | $b_{myc}$ | $3.3\times10^{-5}$ | s.u. $h^{-1}$ |
| MV mediated transcription rate of Myc mRNA | $k_{Mcm1}$ | 0.033 | $h^{-1}$ |
| DE mediated transcription rate of Myc mRNA | $k_{Mcm2}$ | $1.5\times10^{-5}$ | s.u. $h^{-1}$ |
| Hill constant associated with transcription of Myc mRNA, E2F1 mRNA | $k_{mm}$ | 0.33 | s.u. |
| Hill coefficient associated with transcription of Myc mRNA, E2F1 mRNA | *n* | 2 | - |
| Degradation rate of Myc mRNA | $k_{dMcm}$ | 1.38 | $h^{-1}$ |
| Translation rate of Myc protein | $k_{Mcp}$ | 40 | $h^{-1}$ |
| Degradation rate of Myc protein | $k_{dMcp}$ | 0.7 | $h^{-1}$ |
| Skp2 mediated degradation rate of Myc protein | $k_{dMcp1}$ | 2 | $h^{-1}$ |
| Michaelis-Menten constant associated with Skp2 mediated Myc degradation | $J_{MS}$ | 0.001 | s.u. |
| Basal transcription rate of E2F1 mRNA | $b_{E2F}$ | $3.3\times10^{-6}$ | s.u. $h^{-1}$ |
| Myc mediated transcription rate of E2F1 mRNA | $k_{Em1}$ | 0.005 | $h^{-1}$ |
| DE Mediated transcription rate of E2F1 mRNA | $k_{Em2}$ | 0.0005 | s.u. $h^{-1}$ |
| Degradation rate of E2F1 mRNA | $k_{dEm}$ | 0.25 | $h^{-1}$ |
| Translation rate of E2F1 protein | $k_{Ep1}$ | 50 | $h^{-1}$ |
| Degradation rate of E2F1 protein, DE, IDE | $k_{dEp}$ | 0.25 | $h^{-1}$ |
| Cyclin A mediated degradation rate of E2F1 protein | $k_{dEp1}$ | 1 | $h^{-1}$ |
| Michaelis-Menten constant associated with Cyclin A mediated DE degradation | $J_{DE}$ | 1 | s.u. |
| Dp1 and E2F1 association constant | $k_{DE1}$ | 871.4 | s.u.$^{-1}$ $h^{-1}$ |
| DE dissociation constant | $k_{DE2}$ | 55 | $h^{-1}$ |
| Total Dp1 protein | $Dp1_t$ | 1 | s.u. |
| Transcription rate of Rb mRNA | $k_{Rbm}$ | 0.05 | s.u. $h^{-1}$ |
| Degradation rate of Rb mRNA | $k_{dRbm}$ | 1.04 | $h^{-1}$ |
| Translation rate of Rb protein | $k_{Rbp}$ | 5 | $h^{-1}$ |
| Degradation rate of Rb, Rbp, Rbpp protein | $k_{dRbp}$ | 0.231 | $h^{-1}$ |
| DE and Rb association constant | $k_{RD1}$ | 100 | s.u.$^{-1}$ $h^{-1}$ |
| IDE dissociation constant | $k_{RD2}$ | 0.5 | $h^{-1}$ |
| DE and Rbp association constant | $k_{RpD1}$ | 0.8 | s.u.$^{-1}$ $h^{-1}$ |
| IDEP dissociation constant | $k_{RpD2}$ | 0.01 | $h^{-1}$ |

| | | | |
|---|---|---|---|
| Cyclin D mediated phosphorylation rate of free Rb and Rb bound to DE | $k_{iR1}$ | 0.4 | $h^{-1}$ |
| Michaelis-Menten constant associated with Cyclin D mediated phosphorylation of free Rb and Rb bound to DE | $J_{RD}$ | 0.02 | s.u. |
| Dephosphorylation rate of free Rbp and Rbp bound to DE | $k_{iR2}$ | 1.1 | s.u. $h^{-1}$ |
| Michaelis-Menten constant associated with dephosphorylation of free Rbp and Rbp bound to DE | $J_{R1}$ | 0.01 | s.u. |
| Cyclin E mediated phosphorylation rate of free Rbp and Rbp bound to DE | $k_{iR3}$ | 70 | $h^{-1}$ |
| Michaelis-Menten constant associated with Cyclin E mediated phosphorylation of free Rbp and Rbp bound to DE | $J_{RE}$ | 0.005 | s.u. |
| Dephosphorylation rate of free Rbpp | $k_{iR4}$ | 25 | s.u. $h^{-1}$ |
| Michaelis-Menten constant associated with dephosphorylation of free Rbpp | $J_{R2}$ | 0.01 | s.u. |
| MV mediated transcription rate of Cyclin D mRNA | $k_{CDm1}$ | $1.0\times10^{-6}$ | $h^{-1}$ |
| Myc mediated transcription rate of Cyclin D mRNA | $k_{CDm2}$ | 0.01 | $h^{-1}$ |
| Degradation rate of Cyclin D mRNA | $k_{dCDm}$ | 0.173 | $h^{-1}$ |
| Translation rate of Cyclin D protein | $k_{CDp1}$ | 56.67 | $h^{-1}$ |
| Degradation rate of Cyclin D protein | $k_{dCDp}$ | 1.386 | $h^{-1}$ |
| Cyclin A mediated degradation rate of Cyclin D protein | $k_{dCDp1}$ | 2 | $h^{-1}$ |
| Michaelis-Menten constant associated with Cyclin A mediated Cyclin D degradation | $J_{CD}$ | 0.01 | s.u. |
| Basal transcription rate of Cyclin E mRNA | $b_{CycEm}$ | $2.0\times10^{-6}$ | s.u. $h^{-1}$ |
| DE mediated transcription rate of Cyclin E mRNA | $k_{sem}$ | 0.05 | $h^{-1}$ |
| Degradation rate of Cyclin E mRNA | $k_{dcem}$ | 1 | $h^{-1}$ |
| Translation rate of Cyclin E protein | $k_{se}$ | 100 | $h^{-1}$ |
| Degradation rate of Cyclin E protein | $k_{de}$ | 1 | $h^{-1}$ |
| Skp2 mediated degradation rate of Cyclin E protein | $k_{dea}$ | 2 | s.u.$^{-1}$ $h^{-1}$ |
| Transcription rate of p21 mRNA | $k_{sim}$ | 0.0142 | s.u. $h^{-1}$ |
| Degradation rate of p21 mRNA | $k_{dim}$ | 0.693 | $h^{-1}$ |
| Translation rate of p21 protein | $k_{si}$ | 100 | $h^{-1}$ |
| Degradation rate of p21 protein | $k_{di}$ | 1 | $h^{-1}$ |
| Cyclin E and p21 association constant | $k_{aei}$ | 20 | s.u.$^{-1}$ $h^{-1}$ |
| Cyclin E: p21 dissociation constant | $k_{dei}$ | 1 | $h^{-1}$ |
| Cyclin E mediated phosphorylation rate of p21 | $k_{ip}$ | 5 | $h^{-1}$ |
| Michaelis-Menten constant associated with Cyclin E mediated p21 phosphorylation | $J_{ce}$ | 0.1 | s.u. |
| Skp2 mediated degradation rate of p21 protein from Cyclin E: p21 complex | $k_{dip1}$ | 1 | $h^{-1}$ |
| Skp2 mediated degradation rate of phosphorylated p21 | $k_{dip}$ | 10 | s.u.$^{-1}$ $h^{-1}$ |
| Transcription rate of Skp2 mRNA | $k_{17m}$ | 0.0036 | s.u. $h^{-1}$ |
| Degradation rate of Skp2 mRNA | $k_{17dm}$ | 0.17 | $h^{-1}$ |
| Translation rate of Skp2 protein | $k_{17}$ | 26.471 | $h^{-1}$ |
| Degradation rate of dephosphorylated Skp2 protein | $k_{18a}$ | 19.412 | $h^{-1}$ |
| Cdh1 mediated degradation rate of dephosphorylated Skp2 protein | $k_{18b}$ | 52.941 | $h^{-1}$ |
| Michaelis-Menten constant associated with Cdh1 mediated dephosphorylated Skp2 degradation | $J_{18}$ | 0.001 | s.u. |
| Cyclin E mediated phosphorylation rate of Skp2 | $k_{19a}$ | 70.588 | s.u.$^{-1}$ $h^{-1}$ |
| Dephosphorylation rate of Skp2 | $k_{19b}$ | 0.35294 | $h^{-1}$ |
| Degradation rate of phosphorylated Skp2 | $k_{18c}$ | 0.028235 | $h^{-1}$ |
| Cdh1 mediated degradation rate of phosphorylated Skp2 | $k_{18d}$ | 0.35294 | $h^{-1}$ |
| Michaelis-Menten constant associated with Cdh1 mediated phosphorylated Skp2 degradation | $J_{18b}$ | 0.01 | s.u. |
| Basal transcription rate of Cyclin A mRNA | $b_{sam}$ | $1.0\times10^{-6}$ | s.u. $h^{-1}$ |
| DE mediated transcription rate of Cyclin A mRNA | $k_{sam}$ | 0.01 | $h^{-1}$ |

| | | | |
|---|---|---|---|
| Degradation rate of Cyclin A mRNA | $k_{dam}$ | 1 | $h^{-1}$ |
| Translation rate of Cyclin A protein | $k_{sa}$ | 7.0588 | $h^{-1}$ |
| Degradation rate of Cyclin A protein | $k_{da}$ | 0.14118 | $h^{-1}$ |
| Cdh1 mediated degradation rate of Cyclin A protein | $k_{da1}$ | 3.5294 | $s.u.^{-1}\ h^{-1}$ |
| Transcription rate of Cdh1 mRNA | $k_{3m}$ | 0.06 | $s.u.\ h^{-1}$ |
| Degradation rate of Cdh1 mRNA | $k_{3dm}$ | 3 | $h^{-1}$ |
| Translation rate of Cdh1protein | $k_3$ | 176.47 | $h^{-1}$ |
| Degradation rate of Cdh1 protein | $k_{3d}$ | 3.5294 | $h^{-1}$ |
| Basal dephosphorylation rate of Cdh1 protein | $k_{3a}$ | 3.5294 | $s.u.\ h^{-1}$ |
| Cdc20 mediated dephosphorylation rate of Cdh1 protein | $k_{3b}$ | 176.47 | $h^{-1}$ |
| Michaelis-Menten constant associated with Cdc20 mediated Cdh1 dephosphorylation | $J_3$ | 0.1 | s.u. |
| Cyclin B mediated phosphorylation rate of Cdh1 | $k_4$ | 123.53 | $h^{-1}$ |
| Cyclin A mediated phosphorylation rate of Cdh1 | $k_{4b}$ | 123.53 | $h^{-1}$ |
| Michaelis-Menten constant associated with Cyclin A mediated Cdh1 phosphorylation | $J_4$ | 0.04 | s.u. |
| Basal synthesis rate of Cdc20 mRNA | $k_{5am}$ | 0.00035294 | $s.u.\ h^{-1}$ |
| Cyclin B mediated transcription rate of Cdc20 mRNA | $k_{5bm}$ | 0.014118 | $s.u.\ h^{-1}$ |
| Hill constant associated with Cyclin B mediated Cdc20 transcription | $J_5$ | 0.4 | s.u. |
| Hill coefficient associated with Cyclin B mediated Cdc20 transcription | $m$ | 4 | - |
| Degradation rate of Cdc20 mRNA | $k_{5dm}$ | 0.35294 | $h^{-1}$ |
| Translation rate of Cdc20 protein | $k_{5a}$ | 17.647 | $h^{-1}$ |
| Degradation rate of Cdc20 protein | $k_6$ | 0.35294 | $h^{-1}$ |
| IEP mediated activation rate of Cdc20 protein | $k7$ | 2.0 | $h^{-1}$ |
| Michaelis-Menten constant associated with IEP mediated activation of Cdc20 protein | $J_7$ | 0.05 | s.u. |
| Mad mediated deactivation rate of Cdc20 protein | $k_8$ | 1.7647 | $h^{-1}$ |
| Michaelis-Menten constant associated with Mad mediated deactivation of Cdc20 protein | $J_8$ | 0.05 | s.u. |
| Cyclin B mediated phosphorylation rate of IEP | $k_9$ | 0.35294 | $s.u.^{-1}\ h^{-1}$ |
| Dephosphorylation rate of IEP | $k_{10}$ | 0.070588 | $h^{-1}$ |
| Total Mad1 protein | $Mad$ | 1 | s.u. |
| Total IEP protein | $IEP_T$ | 1 | s.u. |
| Transcription rate of Cdt1 mRNA | $k_{21m}$ | 0.0075 | $s.u.\ h^{-1}$ |
| Degradation rate of Cdt1 mRNA | $k_{21dm}$ | 0.35 | $h^{-1}$ |
| Translation rate of Cdt1 protein | $k_{21}$ | 8.8235 | $h^{-1}$ |
| Degradation rate of Cdt1 protein | $k_{22a}$ | 0.21176 | $h^{-1}$ |
| Skp2 mediated degradation rate of Cdt1protein | $k_{22b}$ | 7.0588 | $h^{-1}$ |
| Michaelis-Menten constant associated with Skp2 mediated degradation of Cdt1 protein | $J_{22}$ | 0.04 | s.u. |
| Transcription rate of Geminin mRNA | $k_{20m}$ | 0.0075 | $s.u.\ h^{-1}$ |
| Degradation rate of Geminin mRNA | $k_{20dm}$ | 0.35 | $h^{-1}$ |
| Translation rate of Geminin protein | $k_{19}$ | 8.8235 | $h^{-1}$ |
| Degradation rate of Geminin protein | $k_{20a}$ | 0.21176 | $h^{-1}$ |
| Cdh1 mediated degradation rate of Geminin protein | $k_{20b}$ | 3.5294 | $h^{-1}$ |
| Michaelis-Menten constant associated with Cdh1 mediated degradation of Geminin protein | $J_{20}$ | 0.5 | s.u. |
| Cyclin A mediated transcription rate of Cyclin B mRNA | $k_{scbm}$ | 0.02 | $h^{-1}$ |
| Degradation rate of Cyclin B mRNA | $k_{dcbm}$ | 0.1 | $h^{-1}$ |
| Translation rate of Cyclin B protein | $k_{scb}$ | 3.5294 | $h^{-1}$ |
| Degradation rate of Cyclin B protein | $k_{dcb}$ | 0.14118 | $h^{-1}$ |
| Cdh1 mediated degradation rate of Cyclin B protein | $k_{dcb1}$ | 3.5294 | $s.u.^{-1}\ h^{-1}$ |
| Cdc25 mediate phosphorylation rate of Cyclin B | $k_{cba}$ | 35.294 | $s.u.^{-1}\ h^{-1}$ |
| Wee1 mediated dephosphorylation rate of Cyclin B | $k_{cbi}$ | 35.294 | $s.u.^{-1}\ h^{-1}$ |

| | | | |
|---|---|---|---|
| Cyclin A mediated transcription rate of Cdc25 mRNA | $k_{sc25m}$ | 0.02 | $h^{-1}$ |
| Degradation rate of Cdc25 mRNA | $k_{dc25m}$ | 1 | $h^{-1}$ |
| Translation rate of Cdc25 protein | $k_{sc25}$ | 176.47 | $h^{-1}$ |
| Degradation rate of Cdc25 protein | $k_{dc25}$ | 3.5294 | $h^{-1}$ |
| Cyclin B mediate phosphorylation rate of Cdc25 | $k_{c25a}$ | 35.294 | $h^{-1}$ |
| Dephosphorylation rate of Cdc25 | $k_{c25i}$ | 1.4118 | s.u. $h^{-1}$ |
| Michaelis-Menten constant associated with Cyclin B mediate phosphorylation of Cdc25 | $k_{mc1}$ | 0.05 | s.u. |
| Michaelis-Menten constant associated with dephosphorylation of Cdc25 | $k_{mc2}$ | 0.05 | s.u. |
| Transcription rate of Wee1 mRNA | $k_{swm}$ | 0.016 | s.u. $h^{-1}$ |
| Michaelis-Menten constant associated with Wee1 mRNA transcription | $sgw$ | 1 | - |
| General serum mediated activation rate of Wee1 transcription | $kgw$ | 0.5 | - |
| Degradation rate of Wee1 mRNA | $k_{dwm}$ | 1 | $h^{-1}$ |
| Translation rate of Wee1 protein | $k_{sw}$ | 176.47 | $h^{-1}$ |
| Degradation rate of Wee1 protein | $k_{dw}$ | 3.5294 | $h^{-1}$ |
| Dephosphorylation rate of Wee1 | $k_{wa}$ | 14.118 | s.u. $h^{-1}$ |
| Cyclin B mediate phosphorylation rate of Wee1 | $k_{wi}$ | 176.47 | $h^{-1}$ |
| Michaelis-Menten constant associated with dephosphorylation of Wee1 | $k_{mw1}$ | 0.2 | s.u. |
| Michaelis-Menten constant associated with Cyclin B mediate phosphorylation of Wee1 | $k_{mw2}$ | 0.2 | s.u. |
| p53 dimer mediated activation of p21 transcription | $kp53d$ | 0.0002 | $h^{-1}$ |
| p53 tertramer mediated activation of p21 transcription | $kp53t$ | 0.00078 | $h^{-1}$ |
| Scaling constant to modify cell cycle oscillation period (multiplied with all cell cycle module equations) | $cs$ | 0.51 | |

**Table- S7** Abbreviated forms and description of variables in cell cycle regulatory network **(Govindaraj et al., 2022 ACS Synth. Biol.)**

| Abbreviated form | Description |
|---|---|
| $MV$ | Upstream signalling protein |
| $Myc_m$ | Myc mRNA |
| $Myc_p$ | Myc protein |
| $E2F_m$ | E2F mRNA |
| $E2F_p$ | E2F protein |
| $DE$ | DE complex |
| $iDE$ | iDE complex |
| $iDEP$ | iDEP complex |
| $Rb_m$ | Rb mRNA |
| $Rb$ | Rb protein |
| $CycD_m$ | Cyclin D mRNA |
| $CycD$ | Cyclin D |
| $CycE_m$ | Cyclin E mRNA |
| $CycE$ | Cyclin E |
| $I_m$ | p21 mRNA |
| $I$ | p21 protein |
| $EI$ | Cyclin E-p21 complex |
| $I_p$ | p21 Phosphorylated form |
| $Skp2_m$ | Skp2 mRNA |
| $Skp2_i$ | Skp2 protein |
| $Cdh1_m$ | Cdh1 mRNA |

| | |
|---|---|
| $Cdh1_i$ | Cdh1 protein |
| $CycA_m$ | Cyclin A mRNA |
| $CycA$ | Cyclin A protein |
| $CycB_m$ | Cyclin B mRNA |
| $CycB_a$ | Cyclin B protein |
| $Wee1_m$ | Wee1 mRNA |
| $Wee1_a$ | Wee1 protein |
| $Cdc25_m$ | Cdc25 mRNA |
| $Cdc25_i$ | Cdc25 protein |
| $Cdc20_m$ | Cdc20 mRNA protein |
| $Cdc20_i$ | Cdc20 protein |
| $IEP_a$ | IEP protein |
| $Cdt1_m$ | Cdt1 mRNA |
| $Cdt1$ | Cdt1 mRNA |
| $Gem_m$ | Geminin mRNA |
| $Gem$ | Geminin |

**Table- S8** Reaction governing the proposed cell cycle network **(Govindaraj et al., 2022 ACS Synth. Biol.)**

| S.No | Reaction | Kinetics | Description |
|---|---|---|---|
| **MV variable** | | | |
| R1 | $\xrightarrow{Serum} MV$ | $\frac{k_{mV} \times s \times GF}{smv + GF}$ | Serum mediated synthesis of MV |
| R2 | $MV \rightarrow ::$ | $k_{dmv} \times MV$ | Degradation of MV |
| **Myc mRNA** | | | |
| R3 | $\rightarrow Myc_m$ | $b_{myc} \times s$ | Basal synthesis of Myc mRNA |
| R4 | $\xrightarrow{MV} Myc_m$ | $k_{Mcm1} \times MV$ | MV mediated synthesis of Myc mRNA |
| R5 | $\xrightarrow{DE} Myc_m$ | $\frac{k_{Mcm2} \times s \times DE^n}{(k_{mm}^n \times s^n) + DE^n}$ | DE mediated synthesis of Myc mRNA |
| R6 | $Myc_m \rightarrow ::$ | $k_{dMcm} \times Myc_m$ | Degradation of Myc mRNA |
| **Myc protein** | | | |
| R7 | $\xrightarrow{Myc_m} Myc_p$ | $k_{Mcp} \times Myc_m$ | Synthesis of Myc protein |
| R8 | $Myc_p \rightarrow ::$ | $k_{dMcp} \times Myc_p$ | Degradation of Myc protein |
| R9 | $Myc_p \xrightarrow{Skp2_a} ::$ | $\frac{k_{dMcp1} \times Myc_p \times Skp2}{(s \times J_{ms}) + Myc_p}$ | Skp2 mediated degradation of Myc protein |
| **E2F mRNA** | | | |
| R10 | $\rightarrow E2F_m$ | $b_{e2f} \times s$ | Basal synthesis of E2F mRNA |
| R11 | $\xrightarrow{Myc_p} E2F_m$ | $k_{Em1} \times Myc_p$ | Myc mediated synthesis of E2F mRNA |
| R12 | $\xrightarrow{DE} E2F_m$ | $\frac{k_{Em2} \times s \times DE^n}{(k_{mm}^n \times s^n) + DE^n}$ | DE mediated synthesis of E2F mRNA |

| | | | |
|---|---|---|---|
| R13 | $E2F_m \rightarrow ::$ | $k_{dEm} \times E2F_m$ | Degradation of E2F mRNA |
| **E2F protein** | | | |
| R14 | $\xrightarrow{E2F_m} E2F_p$ | $k_{Ep1} \times E2F_m$ | Synthesis of E2F protein |
| R15 | $E2F_p \rightarrow ::$ | $k_{dEp} \times E2F_p$ | Degradation of E2F protein |
| R16 | $E2F_p \xrightarrow{CycA} ::$ | $\frac{k_{dEp1} \times CycA \times E2F_p}{(s \times J_{DE}) + E2F_p}$ | CycA mediated degradation of E2F protein |
| **DE complex** | | | |
| R17 | $E2F_p + Dp1_p \rightarrow DE$ | $\frac{k_{DE1}}{s} \times Dp1_p \times E2F_p$ | Complex formation of Dp1 and E2F protein |
| R18 | $DE \rightarrow E2F_p + Dp1_p$ | $k_{DE2} \times DE$ | Dissociation of DE complex into Dp1 and E2F protein |
| R19 | $DE \rightarrow Dp1_p + ::$ | $k_{dEp} \times DE$ | Degradation of DE protein |
| R20 | $DE \xrightarrow{CycA} Dp1_p + ::$ | $\frac{k_{dEp1} \times CycA \times DE}{(s \times J_{DE}) + DE}$ | CycA mediated degradation of DE protein |
| **iDE complex** | | | |
| R21 | $DE + Rb \rightarrow iDE$ | $\frac{k_{RD1}}{s} \times DE \times Rb$ | Complex formation of Rb and DE protein |
| R22 | $iDE \rightarrow DE + Rb$ | $k_{RD2} \times iDE$ | Dissociation of inactive DE complex into Rb and E2F protein |
| R23 | $iDE \rightarrow Dp1_p + ::$ | $k_{dEp} \times iDE$ | Degradation of inactive DE protein |
| **iDEP complex** | | | |
| R24 | $DE + iRb1 \rightarrow iDEP$ | $\frac{k_{RpD1}}{s} \times DE \times iRb1$ | Complex formation of single phosphorylated Rb and DE complex |
| R25 | $iDEP \rightarrow DE + iRb1$ | $k_{RpD2} \times iDEP$ | Dissociation of iDEP complex into single phosphorylated Rb and DE complex |
| R26 | $iDE \xrightarrow{CycD} iDEP$ | $\frac{k_{iR1} \times iDE \times CycD}{(J_{RD} \times s) + iDE}$ | First phosphorylation of Rb in iDE by cyclin D |
| R27 | $iDEP \longrightarrow iDE$ | $\frac{k_{iR2} \times s \times iDEP}{(J_{R1} \times s) + iDEP}$ | Dephosphorylation of single phosphorylated Rb in iDEP |
| R28 | $iDEP \xrightarrow{CycE} iRb2 + DE$ | $\frac{k_{iR3} \times iDEP \times CycE}{(J_{RE} \times s) + iDEP}$ | Second phosphorylation of Rb in iDEP by cyclin E |
| **Rb mRNA** | | | |
| R29 | $\rightarrow Rb_m$ | $k_{Rbm} \times s$ | Synthesis of Rb mRNA |
| R30 | $Rb_m \rightarrow ::$ | $k_{dRbm} \times Rb_m$ | Degradation of Rb mRNA |
| **Rb protein** | | | |

| | | | |
|---|---|---|---|
| R31 | $\xrightarrow{Rb_m} Rb$ | $k_{Rbp} \times Rb_m$ | Synthesis of Rb protein |
| R32 | $Rb \rightarrow ::$ | $k_{dRbp} \times Rb$ | Degradation of Rb protein |
| R33 | $Rb \xrightarrow{CycD} iRb1$ | $\frac{k_{iR1} \times Rb \times CycD}{(J_{RD} \times s) + Rb}$ | First phosphorylation of Rb by cyclin D |
| R34 | $iRb1 \longrightarrow Rb$ | $\frac{k_{iR2} \times s \times iRb1}{(J_{R1} \times s) + iRb1}$ | Dephosphorylation of single phosphorylated Rb |
| R35 | $iRb1 \xrightarrow{CycE} iRb2$ | $\frac{k_{iR3} \times iRb1 \times CycE}{(J_{RE} \times s) + iRb1}$ | Second phosphorylation of Rb by cyclin E |
| R36 | $iRb2 \longrightarrow iRb1$ | $\frac{k_{iR4} \times s \times iRb2}{(J_{R2} \times s) + iRb2}$ | Dephosphorylation of double phosphorylated Rb |
| R37 | $iRb1 \rightarrow ::$ | $k_{dRbp} \times iRb1$ | Degradation of single phosphorylated Rb |
| R38 | $iRb2 \rightarrow ::$ | $k_{dRbp} \times iRb2$ | Degradation of double phosphorylated Rb |
| **Cyclin D mRNA** | | | |
| R39 | $\xrightarrow{MV} CycD_m$ | $k_{CDm1} \times MV$ | MV mediated synthesis of Cyclin D mRNA |
| R40 | $\xrightarrow{Myc_p} CycD_m$ | $k_{CDm2} \times Myc_p$ | Myc mediated synthesis of Cyclin D mRNA |
| R41 | $CycD_m \rightarrow ::$ | $k_{dCDm} \times CycD_m$ | Degradation of Cyclin D mRNA |
| **Cyclin D** | | | |
| R42 | $\xrightarrow{CycD_m} CycD$ | $k_{CDp1} \times CycD_m$ | Synthesis of Cyclin D protein |
| R43 | $CycD \rightarrow ::$ | $k_{dCDp} \times CycD$ | Degradation of Cyclin D protein |
| R44 | $CycD \xrightarrow{CycA} ::$ | $\frac{k_{dCDp1} \times CycD \times CycA}{(s \times J_{CD}) + CycD}$ | Cyclin A mediated degradation of Cyclin D protein |
| **Cyclin E mRNA** | | | |
| R45 | $\rightarrow CycE_m$ | $b_{Cycem} \times s$ | Synthesis of Cyclin E mRNA |
| R46 | $\xrightarrow{DE} CycE_m$ | $k_{sem} \times DE$ | E2F mediated synthesis of Cyclin E mRNA |
| R47 | $CycE_m \rightarrow ::$ | $k_{dCEm} \times CycE_m$ | Degradation of Cyclin E mRNA |
| **Cyclin E** | | | |
| R48 | $\xrightarrow{CycE_m} CycE$ | $k_{se} \times CycE_m$ | Synthesis of Cyclin E protein |
| R49 | $CycE \rightarrow ::$ | $k_{dE} \times CycE$ | Degradation of Cyclin E protein |
| R50 | $CycE \xrightarrow{Skp2_a} ::$ | $\frac{k_{dEA}}{s} \times CycE \times Skp2_a$ | Skp2 mediated degradation of Cyclin E protein |
| **p21 mRNA** | | | |
| R51 | $\rightarrow I_m$ | $k_{sim} \times s$ | Synthesis of p21 mRNA |

| | | | |
|---|---|---|---|
| R52 | $I_m \rightarrow ::$ | $k_{dim} \times I_m$ | Degradation of p21 mRNA |
| **p21 protein** | | | |
| R53 | $\xrightarrow{I_m} I$ | $k_{si} \times I_m$ | Synthesis of p21 protein |
| R54 | $I \rightarrow ::$ | $k_{di} \times I$ | Degradation of p21 protein |
| **Cyclin E-p21 complex** | | | |
| R55 | $CycE + I \rightarrow EI$ | $\frac{k_{aei}}{s} \times CycE \times I$ | Complex formation of Cyclin E and p21 |
| R56 | $EI \rightarrow CycE + I$ | $k_{dei} \times EI$ | Dissociation of Cyclin E: p21 complex |
| R57 | $EI \rightarrow ::$ | $k_{di} \times EI$ | Degradation of Cyclin E: p21 complex |
| R58 | $EI \rightarrow ::$ | $k_{dE} \times EI$ | Degradation of Cyclin E: p21 complex |
| R59 | $EI \xrightarrow{Skp2_a} CycE + ::$ | $\frac{k_{dip}}{s} \times EI \times Skp2_a$ | Skp2 mediated degradation of I in EI complex to give free CycE |
| **p21 Phosphorylated form** | | | |
| R60 | $I \xrightarrow{CycE} I_p$ | $\frac{k_{ip} \times CycE \times I}{(J_{CE} \times s) + I}$ | Cyclin E mediated phosphorylation of p21 |
| R61 | $I_p \rightarrow ::$ | $k_{dip1} \times I_p$ | Degradation of phosphorylated p21 |
| R62 | $I_p \xrightarrow{Skp2_a} ::$ | $\frac{k_{dip}}{s} \times I_p \times Skp2_a$ | SKp2 mediated degradation of phosphorylated p21 |
| **Skp2 mRNA** | | | |
| R63 | $\rightarrow Skp2_m$ | $k_{17m} \times s$ | Synthesis of Skp2 mRNA |
| R64 | $Skp2_m \rightarrow ::$ | $k_{17dm} \times Skp2_m$ | Degradation of Skp2 mRNA |
| **Skp2 protein** | | | |
| R65 | $\xrightarrow{Skp2_m} Skp2_i$ | $k_{17} \times Skp2_m$ | Synthesis of Skp2 protein |
| R66 | $Skp2_i \rightarrow ::$ | $k_{18a} \times (Skp2_T - Skp2_a)$ | Degradation of inactive form of Skp2 protein |
| R67 | $Skp2_i \xrightarrow{Cdh1_a} ::$ | $\frac{k_{18b} \times (Skp2_T - Skp2_a)}{(J_{18} \times s) + Skp2_T - \text{[illegible]}}$ | Cdh1 mediated degradation of inactive form of Skp2 protein |
| R68 | $Skp2_i \xrightarrow{CycE} Skp2_a$ | $\frac{k_{19a}}{s} \times CycE \times (Skp2_T - Skp2_a)$ | Cyclin E mediated phosphorylation of Skp2 |

| | | | |
|---|---|---|---|
| R69 | $Skp2_a \rightarrow Skp2_i$ | $k_{19b} \times Skp2_a$ | Dephosphorylation of phosphorylated Skp2 |
| R70 | $Skp2_a \rightarrow ::$ | $k_{18c} \times Skp2_a$ | Degradation of phosphorylated Skp2 protein |
| R71 | $Skp2_a \xrightarrow{Cdh1_a} ::$ | $\frac{k_{18d} \times Skp2_a \times Cdh1_a}{(J_{18b} \times s) + Skp2_a}$ | Cdh1 mediated degradation of phosphorylated Skp2 protein |
| **Cdh1 mRNA** | | | |
| R72 | $\rightarrow Cdh1_m$ | $k_{3m} \times s$ | Synthesis of Cdh1 mRNA |
| R73 | $Cdh1_m \rightarrow ::$ | $k_{3dm} \times Cdh1_m$ | Degradation of Cdh1 mRNA |
| **Cdh1 protein** | | | |
| R74 | $\xrightarrow{Cdh1_m} Cdh1_i$ | $k_3 \times Cdh1_m$ | Synthesis of Cdh1 protein |
| R75 | $Cdh1_i \rightarrow ::$ | $(k_{3d} \times Cdh1_T - Cdh1_a)$ | Degradation of inactive form of Cdh1 protein |
| R76 | $Cdh1_i \rightarrow Cdh1_a$ | $\frac{(k_{3a} \times s) \times (Cdh1_T - Cd\text{[illegible]}}{(J_3 \times s) + Cdh1_T - Cd\text{[illegible]}}$ | Basal activation of Cdh1 protein |
| R77 | $Cdh1_i \xrightarrow{Cdc20_a} Cdh1_a$ | $\frac{k_{3b} \times Cdc20_A \times (Cdh1_T\text{[illegible]}}{(J_3 \times s) + Cdh1_T - \text{[illegible]}}$ | Cdc20 mediated activation of Cdh1 protein |
| R78 | $Cdh1_a \xrightarrow{CycB_a} Cdh1_i$ | $\frac{k_4 \times CycB_a \times Cdh1_a}{(J_4 \times s) + Cdh1_a}$ | Cyclin A mediated inactivation of Cdh1 protein |
| R79 | $Cdh1_a \xrightarrow{CycA} Cdh1_i$ | $\frac{k_{4b} \times CycA \times Cdh1_a}{(J_4 \times s) + Cdh1_a}$ | Cyclin E mediated inactivation of Cdh1 protein |
| R80 | $Cdh1_a \rightarrow ::$ | $k_{3d} \times Cdh1_a$ | Degradation of inactive form of Cdh1 protein |
| **Cyclin A mRNA** | | | |
| R81 | $\rightarrow CycA_m$ | $b_{sam}$ | Basal synthesis rate of Cyclin A mRNA |
| R82 | $\xrightarrow{DE} CycA_m$ | $k_{sam} \times DE$ | E2F mediated synthesis of Cyclin A mRNA |
| R83 | $CycA_m \rightarrow ::$ | $k_{dam} \times CycA_m$ | Degradation of Cyclin A mRNA |
| **Cyclin A protein** | | | |
| R84 | $\xrightarrow{CycA_m} CycA$ | $k_{sa} \times CycA_m$ | Synthesis of Cyclin A protein |
| R85 | $CycA \rightarrow ::$ | $k_{da} \times CycA$ | Degradation of Cyclin A protein |
| R86 | $CycA \xrightarrow{Cdh1_a} ::$ | $k_{da1} \times CycA \times Cdh1_a$ | Cdh1 mediated Cyclin A protein |
| **Cyclin B mRNA** | | | |
| R87 | $\xrightarrow{CycA} CycB_m$ | $\frac{k_{scbm}}{s} \times CycA$ | Cyclin A mediated synthesis of Cyclin B mRNA |
| R88 | $CycB_m \rightarrow ::$ | $k_{dcbm} \times CycB_m$ | Degradation of Cyclin B mRNA |
| **Cyclin B protein** | | | |

| | | | |
|---|---|---|---|
| R89 | $\xrightarrow{CycB_m} CycB_a$ | $k_{scb} \times CycB_m$ | Synthesis of Cyclin B protein |
| R90 | $CycB_a \rightarrow ::$ | $k_{dcb} \times CycB_a$ | Degradation of Cyclin B protein |
| R91 | $CycB_a \xrightarrow{Cdh1_a} ::$ | $\frac{k_{dcb1}}{s} \times Cdh1 \times CycB_a$ | Cdh1 mediated Cyclin B protein |
| R92 | $CycB_a \xrightarrow{Wee1_a} CycB_i$ | $\frac{k_{cbi}}{s} \times Wee1_a \times CycB_a$ | Wee1 mediated dephosphorylation of Cyclin B protein |
| R93 | $CycB_a \xrightarrow{Cdc25_a} CycB_i$ | $\frac{k_{cba}}{s} \times C25_a \times CycB_i$ | Cdc25 mediated phosphorylation of Cyclin B protein |
| R94 | $CycB_i \rightarrow ::$ | $k_{dcb} \times CycB_i$ | Degradation of inactive form of Cyclin B protein |
| R95 | $CycB_i \xrightarrow{Cdh1_a} ::$ | $\frac{k_{dcb1}}{s} \times Cdh1 \times CycB_i$ | Degradation of active form of Cyclin B protein |
| **Wee1 mRNA** | | | |
| R96 | $\xrightarrow{Serum} Wee1_m$ | $\frac{k_{swm} \times s \times Serum}{sgw + (Serum * kgw)}$ | Synthesis of Wee1 mRNA |
| R97 | $Wee1_m \rightarrow ::$ | $k_{dwm} \times Wee1_m$ | Degradation of Wee1 mRNA |
| **Wee1 protein** | | | |
| R98 | $\xrightarrow{Wee1_m} Wee1_a$ | $k_{sw} \times Wee1_m$ | Serum mediated synthesis of Wee1 protein |
| R99 | $Wee1_a \rightarrow ::$ | $k_{dw} \times Wee1_a$ | Degradation of Wee1 protein |
| R100 | $Wee1_a \xrightarrow{CycB_a} Wee1_i$ | $\frac{k_{wi} \times Wee1_a \times CycB_a}{(k_{mw2} \times s) + Wee1_a}$ | Cyclin B mediated phosphorylation of Wee1 protein |
| R101 | $Wee1_i \rightarrow Wee1_a$ | $\frac{k_{wa} \times s \times Wee1_i}{(k_{mc1} \times s) + Wee1_i}$ | Dephosphorylation of Wee1 protein |
| R102 | $Wee1_i \rightarrow ::$ | $k_{dw} \times Wee1_i$ | Degradation of inactive form of Wee1 protein |
| **Cdc25 mRNA** | | | |
| R103 | $\xrightarrow{CycA} Cdc25_m$ | $k_{sc25m} \times CycA$ | Cyclin A mediated synthesis of Cdc25 mRNA |
| R104 | $Cdc25_m \rightarrow ::$ | $k_{dc25m} \times C25_m$ | Degradation of Cdc25 mRNA |
| **Cdc25 protein** | | | |
| R105 | $\xrightarrow{Cdc25_m} Cdc25_i$ | $k_{sc25} \times C25_m$ | Synthesis of Cdc25 protein |
| R106 | $Cdc25_i \rightarrow ::$ | $k_{dc25} \times C25_i$ | Degradation of Cdc25 protein |
| R107 | $Cdc25_i \xrightarrow{CycB_a} Cdc25_a$ | $\frac{k_{c25a} \times C25_i \times CycB_a}{(k_{mc1} \times s) + C25_i}$ | Cyclin B mediated phosphorylation of Cdc25 protein |
| R108 | $Cdc25_a \rightarrow Cdc25_i$ | $\frac{k_{c25i} \times s \times C25_a}{(k_{mc2} \times s) + C25_a}$ | Dephosphorylation of Cdc25 protein |

| | | | |
|---|---|---|---|
| R109 | $Cdc25_a \rightarrow ::$ | $k_{dc25} \times C25_a$ | Degradation of inactive form of Cdc25 protein |
| **Cdc20 mRNA protein** | | | |
| R110 | $\rightarrow Cdc20_m$ | $k_{5am} \times s$ | Synthesis of Cdc20 mRNA |
| R111 | $\xrightarrow{CycB_a} Cdc20_m$ | $\dfrac{k_{5bm} \times s \times \left(\frac{CycB_a}{J_5 \times s}\right)^m}{1 + \left(\frac{CycB_a}{(J_5 \times s)}\right)^m}$ | MV and Cyclin A mediated synthesis of Cdc20 mRNA |
| R112 | $Cdc20_m \rightarrow ::$ | $k_{5dm} \times Cdc20_m$ | Degradation of Cdc20 mRNA |
| **Cdc20 protein** | | | |
| R113 | $\xrightarrow{Cdc20_m} Cdc20_i$ | $k_{5a} \times Cdc20_m$ | Synthesis of Cdc20 protein |
| R114 | $Cdc20_i \rightarrow ::$ | $k_6 \times (Cdc20_T - Cdc20_a)$ | Degradation of inactive Cdc20 protein |
| R115 | $Cdc20_i \xrightarrow{IEP_a} Cdc20_a$ | $\dfrac{k_7 \times IEP_a \times (Cdc20_T - \text{[illegible]}}{(J_7 \times s) + Cdc20_T - \text{[illegible]}}$ | IEP mediated activation of Cdc20 protein |
| R116 | $Cdc20_a \xrightarrow{Mad} Cdh1_i$ | $\dfrac{k_8 \times Mad \times s \times Cdc20_a}{(J_8 \times s) + Cdc20_a}$ | Mad mediated inactivation of Cdc20 protein |
| R117 | $Cdc20_a \rightarrow ::$ | $k_6 \times Cdc20_a$ | Degradation of active Cdc20 protein |
| **IEP protein** | | | |
| R118 | $IEP_i \xrightarrow{CycB_a} IEP_a$ | $\frac{k_9}{s} \times CycB_a \times ((IEP_T \times s) - IEP_a)$ | Cyclin A mediated Phosphorylation of IEP protein |
| R119 | $IEP_a \rightarrow IEP_i$ | $k_{10} \times IEP_a$ | Dephosphorylation of phosphorylated IEP protein |
| **Cdt1** | | | |
| R120 | $\rightarrow Cdt1_m$ | $k_{21m} \times s$ | Synthesis of Cdt1 mRNA |
| R121 | $Cdt1_m \rightarrow ::$ | $k_{21dm} \times Cdt1_m$ | Degradation of Cdt1 mRNA |
| R122 | $\xrightarrow{Cdt1_m} Cdt1$ | $k_{21} \times Cdt1_m$ | Synthesis of Cdt1 protein |
| R123 | $Cdt1 \rightarrow ::$ | $k_{22a} \times Cdt1$ | Degradation of Cdt1 protein |
| R124 | $Cdt1 \xrightarrow{Skp2_a} ::$ | $\dfrac{k_{22b} \times Cdt1 \times Skp2_a}{(J_{22} \times s) + Cdt1}$ | Skp2 mediated degradation of Cdt1 protein |
| **Geminin** | | | |
| R125 | $\rightarrow Gem_m$ | $k_{20m} \times s$ | Synthesis of Geminin mRNA |
| R126 | $Gem_m \rightarrow ::$ | $k_{20dm} \times Gem_m$ | Degradation of Geminin mRNA |
| R127 | $\xrightarrow{Gem_m} Gem$ | $k_{19} \times Gem_m$ | Synthesis of Geminin protein |
| R128 | $Gem \rightarrow ::$ | $k_{20a} \times Gem$ | Degradation of Geminin protein |

| | | | |
|---|---|---|---|
| R129 | $Gem \xrightarrow{Cdh1_a} ::$ | $\frac{k_{20b} \times Gem \times Cdh1_a}{(J_{20} \times s) + Gem}$ | Cdh1 mediated degradation of Geminin protein |
| **Coupling of p53-cell cycle model** | | | |
| R130 | $\xrightarrow{p53_2} I_m$ | $kp53d \times p53d$ | p53 dimer induced transcription of p21 mRNA |
| R131 | $\xrightarrow{p53_4} I_m$ | $kp53t \times p53t$ | p53 tetramer induced transcription of p21 mRNA |

**Table S9:** ODEs governing the dynamics of the system for the proposed mitotic catastrophe module in **Fig. 6A.**

| | |
|---|---|
| $\frac{dPIDD}{dt} = k_{spi} + k_{pid} \times p53D + k_{pit} \times p53T - k_{dpi} \times PIDD - k_{pact} \times \frac{PIDD}{k_{gem} + PIDD} \times (\frac{1}{1 + kcdtc \times Cdt1}) \times \frac{DNA_{dam}}{1 + DNA_{dam}}$ | 1 |
| $\frac{dPIDDact}{dt} = (k_{pact}) \times \frac{PIDD}{k_{gem} + PIDD} \times (\frac{1}{1 + kcdtc \times Cdt1}) \times \frac{DNA_{dam}}{1 + DNA_{dam}} - k_{dap} \times PIDD_{act} - k_{com} \times \frac{PIDDact}{k_{pdd} + PIDDact} \times \frac{casp2}{kpcc + casp2} + k_{depc} \times PDCa$ | 2 |
| $\frac{dcasp2}{dt} = k_{scasp} - \frac{k_{11s} \times casp2}{J_{11} + +casp2} - \frac{k_{11a} \times casp2 \times CycA}{J_{12} + casp2} - \frac{k_{11b}\, casp2 \times CycBa}{J_{13} + casp2} - \frac{k_{11c} \times casp2 \times AURKB}{J_{14} + +casp2} + \frac{k_{11d} \times (casp2i)}{J_{15} + (casp2i))} - k_{dcasp} \times casp2 - k_{com} \times \frac{PIDDact}{k_{pdd} + PIDDact} \times casp2 + k_{depc} \times PDCa$ | 3 |
| $\frac{dcasp2i}{dt} = \frac{k_{11s} \times casp2}{J_{11} + +casp2} + \frac{k_{11a} \times casp2 \times CycA}{J_{12} + casp2} + \frac{k_{11b}\, casp2 \times CycBa}{J_{13} + casp2} + \frac{k_{11c} \times casp2 \times AURKB}{J_{14} + +casp2} - k_{dcasp} \times casp2i$ | 4 |
| $\frac{dPDCa}{dt} = k_{com} \times \frac{PIDDact}{k_{pdd} + PIDDact} \times \frac{casp2}{kpcc + casp2} - k_{depc} \times PDCa - k_{degpc} \times \frac{PDca}{j_{deg} + PDca}$ | 5 |
| $\frac{dAURKB}{dt} = ksau - kdau \times AURK - kdauc \times AURKB \times \frac{Cdh1}{jau + AURKB}$ | 6 |
| $\frac{dMdm}{dt} = k_{smdm} + k_{mono} \times P53_{mono} + k_{dimer} \times p53_2 + k_{tetramer} \times p53_4 + k_2 \times Mdm^p - (k_{dmdm} + \frac{k_{dnam} \times DNAdam}{k_a + DNAdam}) \times Mdm - k_1 \times Mdm - k_{PIDDosome} \times Mdm \times \frac{PDca^{mn}}{kpidd^{mn} + PDca^{mn}}$ | 7 |

**Table- S10** Description of the model parameters and their values for mitotic catastrophe model in **Fig. 6A.**

| Description | Parameter | Values | Unit |
|---|---|---|---|
| PIDD synthesis rate | $k_{spi}$ | 0.00000001 | s.u. $h^{-1}$ |
| p53 dimer mediated synthesis rate of PIDD | $k_{pid}$ | 0.000001 | $h^{-1}$ |
| p53 tetramer mediated synthesis rate of PIDD | $k_{pit}$ | 1 | $h^{-1}$ |
| Degradation rate of PIDD | $k_{dpi}$ | 1 | $h^{-1}$ |
| Activation rate of PIDD | $k_{pact}$ | 7.4 | s.u. $h^{-1}$ |
| Michaelis Menten constant for Cdt1 inhibited PIDD activation | $k_{gem}$ | 10.0 | s.u. |
| Deactivation rate of PIDDact | $k_{dap}$ | 0.5 | $h^{-1}$ |
| Cdt1 mediated inhibition of PIDD activation | $k_{cdtc}$ | 20.0 | s.u.$^{-1}$ |
| Complexation rate of PIDDact and Casp-2 complex | $k_{com}$ | 0.24 | s.u.$^{1}$ $h^{-1}$ |
| Michaelis Menten constant for complexation of PIDDact and Casp-2 complex | $k_{pdd}$ | 20 | s.u. |
| Dissociation rate of PDca complex | $k_{depc}$ | 0.0000001 | $h^{-1}$ |
| Synthesis rate of Casp-2 | $k_{scasp}$ | 1 | s.u. $h^{-1}$ |
| Phosphorylation rate of Casp-2 | $k_{11s}$ | 0.352 | s.u. $h^{-1}$ |
| Michaelis Menten constant for phosphorylation of Casp-2 | $J_{11}$ | 0.03 | s.u. |
| CycA mediated phosphorylation rate of Casp-2 | $k_{11a}$ | 10 | $h^{-1}$ |
| CycA mediated phosphorylation rate of Casp-2 | $J_{12}$ | 0.03 | s.u. |
| CycB mediated phosphorylation rate of Casp-2 | $k_{11b}$ | 10.0 | $h^{-1}$ |
| Michaelis Menten constant for CycB mediated phosphorylation rate of Casp-2 | $J_{13}$ | 0.03 | s.u. |
| AURKB mediated phosphorylation rate of Casp-2 | $k_{11c}$ | 20.0 | $h^{-1}$ |
| Michaelis Menten constant for AURKB mediated phosphorylation rate of Casp-2 | $J_{14}$ | 0.03 | s.u. |
| Dephosphorylation rate of Casp-2 | $k_{11d}$ | 1.7 | $h^{-1}$ |
| Michaelis Menten constant for dephosphorylation rate of Casp-2 | $J_{15}$ | 0.03 | s.u. |
| Degradation rate of Casp-2 and Casp2i | $k_{dcasp}$ | 1.0 | $h^{-1}$ |
| Degradation rate of PDca complex | $k_{degpc}$ | 0.008 | s.u. $h^{-1}$ |
| PDca complex mediated degradation rate of Mdm2 | $k_{piddosome}$ | 0.9 | $h^{-1}$ |

| | | | |
|---|---|---|---|
| Hill constant for PDca complex mediated degradation of Mdm2 | $K_{pidd}$ | 0.4 | s.u. |
| Hill coefficient for PDca complex mediated degradation of Mdm2 | $mn$ | 4 | - |
| AURKB synthesis rate | $ksau$ | 0.16 | s.u. $h^{-1}$ |
| AURKB degradation rate | $kdau$ | 0.21 | $h^{-1}$ |
| Cdh1 mediated degradation rate of AUKB | $kdauc$ | 3.52 | $h^{-1}$ |
| Michelis-Menten constant for Cdh1 mediated AUKB degradation rate | $jau$ | 0.5 | s.u. |

**Table- S11** Abbreviated forms and description of variables for the proposed mitotic catastrophe module in **Fig. 6A**:

| Abbreviated form | Description |
|---|---|
| $PIDD$ | p53 induced death domain (PIDD) |
| $PIDD_{act}$ | PIDD active form |
| $Casp2$ | Caspase-2 non phosphorylated form |
| $Casp2i$ | Caspase-2 phosphorylated form |
| $PDca$ | PIDD Caspase-2 complex (PIDDosome) |
| AURKB | Aurora Kinase B protein |

**Table- S12** Reaction governing the proposed mitotic catastrophe module in **Fig. 6A.**

| Reaction number | Reaction | Propensity function |
|---|---|---|
| 1 | $\longrightarrow \mathrm{PIDD}$ | $w1 = k_{spi}$ |
| 2 | $\xrightarrow{p53\mathrm{D}} \mathrm{PIDD}$ | $\mathrm{w}2 = k_{pid} \times p53D$ |
| 3 | $\xrightarrow{p53\mathrm{T}} \mathrm{PIDD}$ | $\mathrm{w}3 = k_{pit} \times p53T$ |
| 4 | $\mathrm{PIDD} \longrightarrow$ | $w4 = k_{dpi} \times \mathrm{PIDD}$ |
| 5 | $\mathrm{PIDD} \longrightarrow \mathrm{PIDDact}$ | $w5 = (k_{pact}/cdt1) \times \frac{PIDD}{k_{gem} + PIDD} \times \frac{DNA_{dam}}{1 + DNA_{dam}}$ |
| 6 | $\mathrm{PIDDact} \longrightarrow PIDD$ | $w6 = k_{dap} \times PIDD_{act}$ |

| | | |
|---|---|---|
| **7** | $\text{PIDDact} + \text{Casp2} \rightarrow PDCa$ | $w7 = k_{com} \times \frac{PIDDact}{k_{pdd} + PIDDact} \times \frac{\mathbf{casp2}}{kpcc + casp2}$ |
| **8** | $- \rightarrow Casp2$ | $w8 = k_{scasp}$ |
| **9** | $Casp2 \rightarrow Casp2p$ | $w9 = \frac{k_{11s} \times casp2}{J_{11} + +casp2}$ |
| **10** | $Casp2 \xrightarrow{\text{CycA}} Casp2p$ | $w10 = \frac{k_{11s} \times casp2}{J_{11} + +casp2}$ |
| **11** | $Casp2 \xrightarrow{\text{CycB}} Casp2p$ | $w11 = \frac{k_{11b}\, casp2 \times CycBa}{J_{13} + casp2}$ |
| **12** | $Casp2 \xrightarrow{\text{AURKB}} Casp2p$ | $w12 = \frac{k_{11c} \times casp2 \times AURKB}{J_{14} + +casp2}$ |
| **13** | $Casp2p \xrightarrow{\text{AURKB}} Casp2$ | $w13 = \frac{k_{11d} \times (casp2i)}{J_{15} + (casp2i)}$ |
| **14** | $\text{casp2} \rightarrow$ | $w14 = k_{dcasp} \times casp2$ |
| **15** | $\text{PDca} \longrightarrow \text{PIDDact} + \text{Casp2}$ | $w15 = k_{depc} \times PDCa$ |
| **16** | $PDca \rightarrow$ | $w16 = k_{degpc} \times \frac{PDca}{j_{deg} + PDca}$ |
| **17** | $Mdm2 \xrightarrow{\text{PDca}}$ | $w17 = -k_{PIDDosome} \times Mdm \times \frac{PDca^{mn}}{kpidd^{mn} + PDca^{mn}}$ |
| **18** | $- \rightarrow AURKB$ | $w18 = ksau$ |
| **19** | $AURKB \rightarrow$ | $w19 = kdau \times AURKB$ |
| **20** | $AURKB \xrightarrow{Cdh1}$ | $w20 - kdauc \times AURKB \times \frac{Cdh1}{jau + AURKB}$ |
